# Comparative Metabolomic Analysis of Vaginal Microbiota in Planktonic and Biofilm States Unveils Species-Specific Metabolic Signatures

**DOI:** 10.1101/2025.03.21.644638

**Authors:** Smrutiti Jena, Damilola Lawore, Leopold N Green, Douglas K. Brubaker

**Affiliations:** Weldon School of Biomedical Engineering, Purdue University, West Lafayette IN; Center for Global Health and Disease, Department of Pathology, Case Western Reserve University, Cleveland OH; The Blood Heart Lung Immunology Research Center, Case Western Reserve University, University Hospitals of Cleveland, Cleveland, OH, USA

**Keywords:** Bacterial vaginosis, vaginal microbes, biofilm, metabolomics, metabolic pathway, metabolites, *L. crispatus*, *G. vaginalis*, *L*. *iners*

## Abstract

Bacterial vaginosis (BV) affects approximately 29% of women in the U.S., with higher rates among certain demographics and up to 50% recurrence within a year. Besides complications like increased risk of sexually transmitted infections (STIs), pregnancy-related issues, it can negatively impact psychological well-being, leading to discomfort and reduced quality of life. While previous studies have provided insights into the overall metabolomic profile of healthy and diseased vaginal environments, the elucidation of individual microbial metabolite signatures remains limited. Furthermore, given that biofilms exhibit distinct metabolic requirements compared to planktonic cultures, a differential analysis of metabolites in both growth conditions could reveal potential therapeutic targets. This study presents a comprehensive metabolomic analysis and comparison of significant vaginal microbes including *Lactobacillus crispatus, Gardenerella vaginalis*, and *Lactobacillus iners* in both planktonic and biofilm growth conditions. Our analysis revealed distinct metabolite production and consumption patterns among different microbes and growth modes. In biofilm cultures, metabolite consumption is influenced by nutrient availability, which in turn regulates the profile of produced metabolites. *G. vaginalis* demonstrated the ability to form biofilms in various media types. Limited shared metabolic pathways in both biofilm types of *G. vaginalis*, highlights the unique metabolic processes involved in their formation. Despite *L. crispatus* suspension and biofilm cultures sharing 142 consumed and 104 produced metabolites, the biofilm culture demonstrated a remarkable metabolic shift. While comparing suspension and biofilm cultures of *L. crispatus, L. iners*, and *G. vaginalis,* we found convergence in nutrient utilization, but divergence in metabolic outputs reflecting growth-specific adaptations and underscore the importance of considering the state of existence when studying the vaginal microbiome.

This study provides valuable insights into the growth mode-specific metabolic requirements of key vaginal microbes. The findings underscore the potential for leveraging metabolite-mediated microbial cross-talk as a novel therapeutic approach against BV. This avenue of research warrants further investigation, as it could lead to the development of targeted interventions that modulate the vaginal microbiome through metabolic manipulation, potentially offering more effective and personalized treatments for BV.

## Introduction

The physiological vaginal microbiota was first characterized in 1892 by Albert Döderlein as a homogeneous population, primarily comprising Gram-positive bacilli believed to originate from the gut, now identified as part of the genus Lactobacillus ^1^. The evolution of this distinctive vaginal microbiome is underpinned by two evolutionary hypotheses: the “disease risk hypothesis” and the “obstetric protection hypothesis”. These theories propose that the human vagina is predominantly populated by protective Lactobacillus species due to increased susceptibility to sexually transmitted diseases and higher risks of pregnancy- and childbirth-associated microbial complications ^2^.

Bacterial vaginosis (BV) is a prevalent vaginal disorder characterized by a spectrum of clinical manifestations, ranging from asymptomatic cases to those presenting with unusual odor, discharge, and irritation. BV occurs when the intravaginal microbial community dynamics shift from Lactobacillus *spp.* dominated to other anaerobic bacteria overgrowth like *Gardnerella vaginalis* ^3^. The vaginal microbiome has been classified into five Community State Types (CSTs I-V), each defined by the predominant microorganism shaping the vaginal microenvironment. CSTs I, II, III, and V are primarily dominated by *L. crispatus, L. gasseri, L. iners*, and *L. jensenii*, respectively. In contrast, CST IV represents a polymicrobial community with minimal presence of Lactobacillus spp., often indicative of the microbial environment associated with BV^4^. BV is linked to a higher risk of acquiring sexually transmitted infections (STIs) such as *N. gonorrhoeae, C. trachomatis, T. vaginalis*, herpes simplex virus, human papillomavirus, and human immunodeficiency virus. Additionally, BV is associated with other conditions such as pelvic inflammatory disease, endometritis, chorioamnionitis, and amniotic fluid infection. During pregnancy, BV has been correlated with adverse outcomes, including preterm premature rupture of membranes, preterm labor, and preterm birth ^5^ ^6^ ^7^ ^8^ ^9,10^.

In a healthy vaginal ecosystem, Lactobacillus *spp.* play a crucial role in maintaining microbial balance through various mechanisms. These organisms produce hydrogen peroxide, which effectively eliminates catalase-deficient microbes, thereby conferring a survival advantage to Lactobacilli ^11^. Furthermore, Lactobacillus species generate an array of antimicrobial peptides, including bacteriocins, bacteriocin-like substances, and biosurfactants, which contribute to their protective function ^12–14^. Additionally, they can stimulate the process of autophagy, leading to the degradation of intracellular bacteria, protozoa, and viruses ^12,15^. All Lactobacilli *spp.*, apart from *L. iners*, are significant producers of both D-lactic acid and L-lactic acid. D-lactic acid provides greater protection against vaginal dysbiosis compared to L-lactic acid. Levels of D-lactic acid is highest when *L. crispatus* is the dominant species and lowest in the presence of *L. iners*, *G. vaginalis*, and other organisms associated with BV ^16^.

*L. iners* displays adaptability to dynamic microbiome conditions as it often dominates during transitional phases of the vaginal microbiota and menstruation. ^17^ ^18^. *L. iners* possesses certain genes that imply its potential as an opportunistic pathogen, including inerolysin, a notable cholesterol-dependent cytolysin. Studies indicate, the presence of *L. iners* in the vaginal niche may not serve as a reliable biomarker for vaginal health, unlike other commonly identified Lactobacillus spp. ^19^ ^20^ ^21^. Moreover, in *L. iners*-dominated microbiomes there is a higher likelihood of transitioning towards dysbiosis when compared to *L. crispatus*-dominated microbiomes ^22^.

*G. vaginalis,* formerly recognized as *Haemophilus vaginalis*, was once regarded as the sole causative agent responsible for clinical symptoms used to diagnose BV ^23,24^. However, an updated model has been proposed which considers recent advancements in animal models for BV and emphasizes the interactions among multiple microbes are more pertinent to understanding the condition. ^25^.

Numerous studies suggest that BV is characterized by the presence of a polymicrobial community, devoid of Lactobacilli *spp.*, organized within a biofilm structure. ^26–28^. Biofilms are characterized as microbial populations enclosed within a self-produced matrix, adhering to each other and to surfaces. Distinct regions within the biofilm may harbor genetically identical cells, yet they can display varying patterns of gene expression conferring enhanced tolerance to unfavorable conditions ^29^. Microorganisms within biofilms exhibit distinct behavior compared to planktonic cells, making them more resilient against conventional antimicrobial treatments and enabling evasion of the host’s immune response ^30^ ^31^. According to Swidsinski, this resistance is why biofilms of *G. vaginalis* in women with BV cannot be eradicated using traditional therapies involving moxifloxacin and metronidazole ^32^. Thus, biofilms may contribute to persistent, slow advancing chronic infections. The layering of epithelial cells with multiple bacterial layers observed in cue cells, a hallmark of the Amsel criteria for diagnosing BV, closely resembles what we would expect to find in biofilm formation. For decades we examined cue cells without recognizing them as indicative of biofilm formation until it was confirmed with visualization of polymicrobial biofilm adhering to vaginal epithelial cells in BV using fluorescence in situ hybridization (FISH) ^33^. Numerous studies have solidified the consensus that biofilms in BV are closely linked with *G. vaginalis*, which emerged as the predominant component despite the biofilm containing elevated levels of various bacterial groups ^34^.

Metabolites, small intermediate products of metabolism, facilitate interspecies metabolic collaboration in bacterial communities, exemplified by oxygen gradients in biofilms enabling anaerobic bacteria to thrive alongside aerobes. ^35^; ^36^ ^37^. There are studies that establish cooperative and syntrophic interactions between microbes through metabolic exchanges in a certain niche significantly dictate nutritional quality and habitants of the community ^38^ ^39,40^. When produced by the host, they serve as components of various structures, contribute to defense systems against pathogens, participate in cell signaling pathways, and act as substrates for microorganisms ^41^. Metabolites play a significant role in fostering interkingdom symbiotic relationships, encompassing mutualistic, commensal, and parasitic interactions. Symbioses between microorganisms and multicellular hosts exemplify the critical bidirectional molecular interactions fueled by metabolites, essential for maintaining these symbiotic relationships. Microbial metabolites are sensed through a well-orchestrated cellular process and induce host responses. Detection of metabolites by one type of host cell can serve to communicate with other cell types within the host, facilitating the coordination of host responses both locally and systemically ^42^ ^43^.

Recent technological advancements have enabled the analysis of metabolome across various cell types. We have developed an analytical pipeline for the interpretation of high throughput metabolomic data derived from microorganisms of significance in the vaginal microbiome, with a focus on species associated with the etiology of BV. This study will provide an insight into the differential production or consumption of metabolites across species and discrete metabolic growth conditions like suspension and biofilm cultures.

## Results

### Certain metabolites are uniquely generated or consumed in the suspension culture of vaginal microbes

Our analysis revealed distinct patterns in the production or consumption of specific metabolites among suspension cultures of *L. crispatus, G. vaginalis* and *L. iners*, as illustrated in Fig 1A. In this representation, blue indicates metabolites consumed, while red indicates those produced, with intensity reflecting the extent of production or consumption. For example, Gluconate was produced by *L. crispatus* and consumed by both *G. vaginalis* and *L. iners*. Adenosine, though consumed by all, was notably more heavily utilized by *L. crispatus*. Metabolites such as glutamine, NAD+, uridine monophosphate, 2-phosphoglycerate, and ribose were exclusively produced by both Lactobacillus species. However, there were metabolites like N-formyl methionine, N-propionylmethionine, and N-acetyl methionine that were produced by *L. iners* and consumed by *L. crispatus*. Similarly, cytosine and lactate were produced by *L. crispatus* and consumed by *L. iners*. Histamine was found to be significantly and exclusively produced by *L. iners*. Pyruvate and deoxycholate were notably more abundantly produced by *G. vaginalis* compared to the other two microbes (Fig 1A).

**Fig 1:**
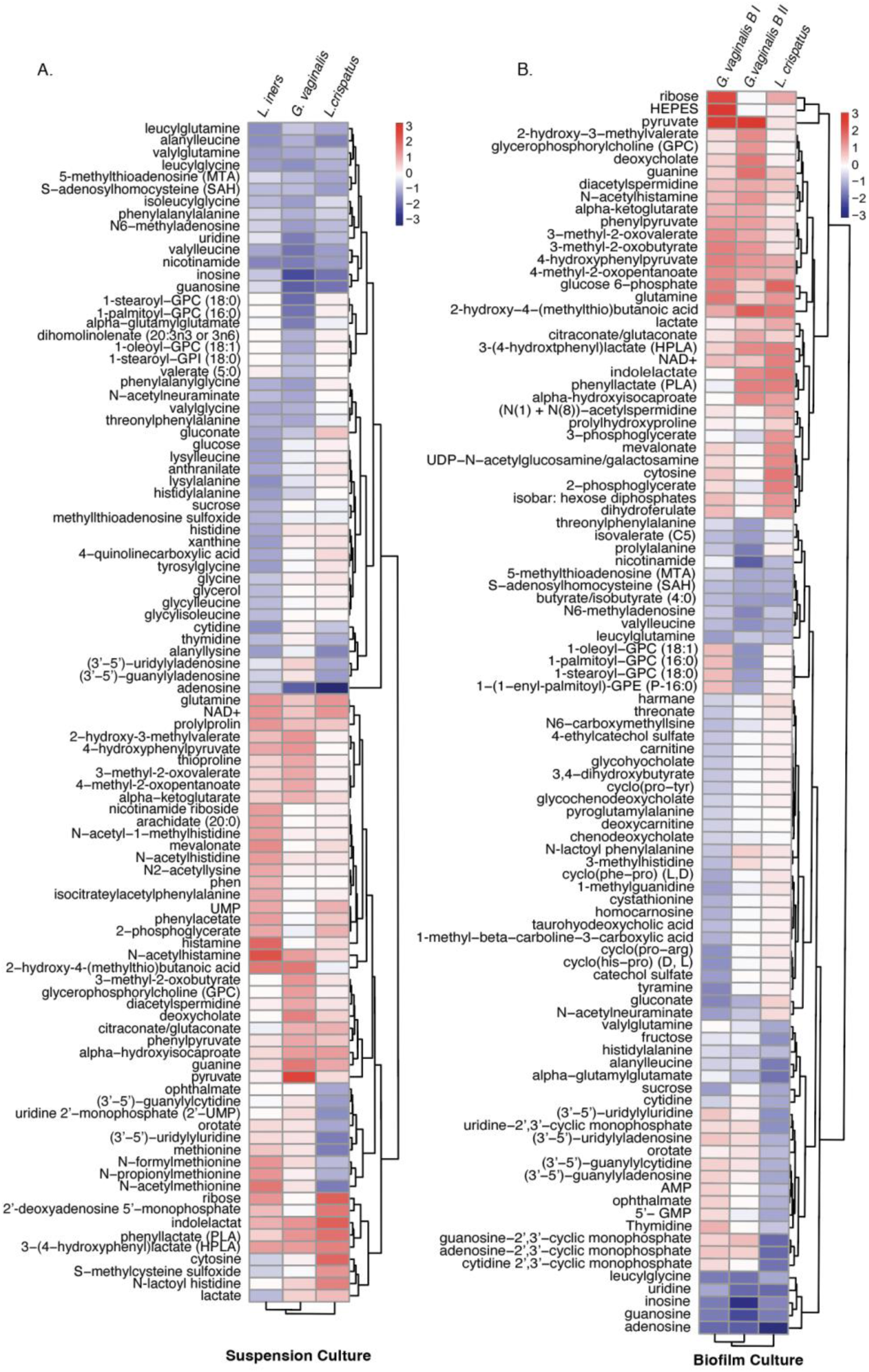
Metabolites generated or consumed by suspension and biofilm cultures across microbes show nutrient-modulated diverse profile. Heatmaps showing differential abundance of metabolites in (A) suspension cultures of *L. crispatus, L. iners* and *G. vaginalis* mapped on log2 (blank media normalized) value and (B) metabolite profile of biofilm culture of *L. crispatus* and both types of *G. vaginalis* biofilms mapped on log2 (blank media normalized) value. *G. vaginalis* B I: Biofilm type I grown in NYCIII media and *G. vaginalis* B II: Biofilm type II grown in supplemented BHI media (sBHI). Red represents produced metabolites, while blue denotes consumed metabolites.

### Consumption of metabolite depends on the substrate availability that regulates the composition of produced metabolites in biofilm cultures

Even though the media (MRS) used for both suspension and biofilm cultures of *L. crispatus* remained consistent, noticeable differences were observed in the consumption and production patterns of metabolites across these cultures. Similar distinctions were evident in the suspension and biofilm type I of *G. vaginalis*. Furthermore, the metabolite profiles differed between type I and type II biofilms of *G. vaginalis*, indicating distinct utilization and production of metabolites in biofilm culture modes, influenced by available substrates (Fig 1B).

*L. crispatus* was the exclusive producer of gluconate, N-acetylneuraminate, and harmane, while both types of *G. vaginalis* biofilms consumed them. Pyruvate, cGMP, and cAMP were solely produced by *G. vaginalis* biofilms, with the latter two being consumed by *L. crispatus*. Notably, in the case of *G. vaginalis* biofilm type II, 1-oleoyl GPC, 1-palmitoyl GPC, 1-stearoyl GPC, and 1−(1−enyl-palmitoyl)-GPE were produced, but in type I, they were consumed. Ribose and HEPES were significantly produced by type II biofilms but not by type I (Fig 1B).

### Enriched metabolite pathways identified from *G. vaginalis* cultures reveal greater similarity between suspension and biofilm type I, while highlighting a significant metabolic distinction between biofilm types I and II

A total of 108 metabolites were commonly consumed, while 32 were produced across all forms of *G. vaginalis* culture (Fig. 2A & 2D). The commonly consumed metabolites across all growth modes were associated with 18 significant metabolic pathways, with phosphatidylcholine biosynthesis emerging as the most prominent, followed closely by the cardiolipin biosynthesis pathway. (Fig. 2B). The 25 metabolites, specifically consumed by suspension culture but none of the biofilm types belonged only either to bile acid biosynthesis or thiamine metabolism (Data not shown). Eleven metabolites were commonly consumed by both biofilm types, primarily associated with taurine and hypotaurine metabolism, followed by homocysteine degradation (Fig 2C). This minimal overlap highlights the metabolic adaptability and diversity of *G. vaginalis* during biofilm formation in response to nutrient availability, demonstrating that the organism can adopt entirely distinct metabolic pathways based on the resources at its disposal.

**Fig 2:**
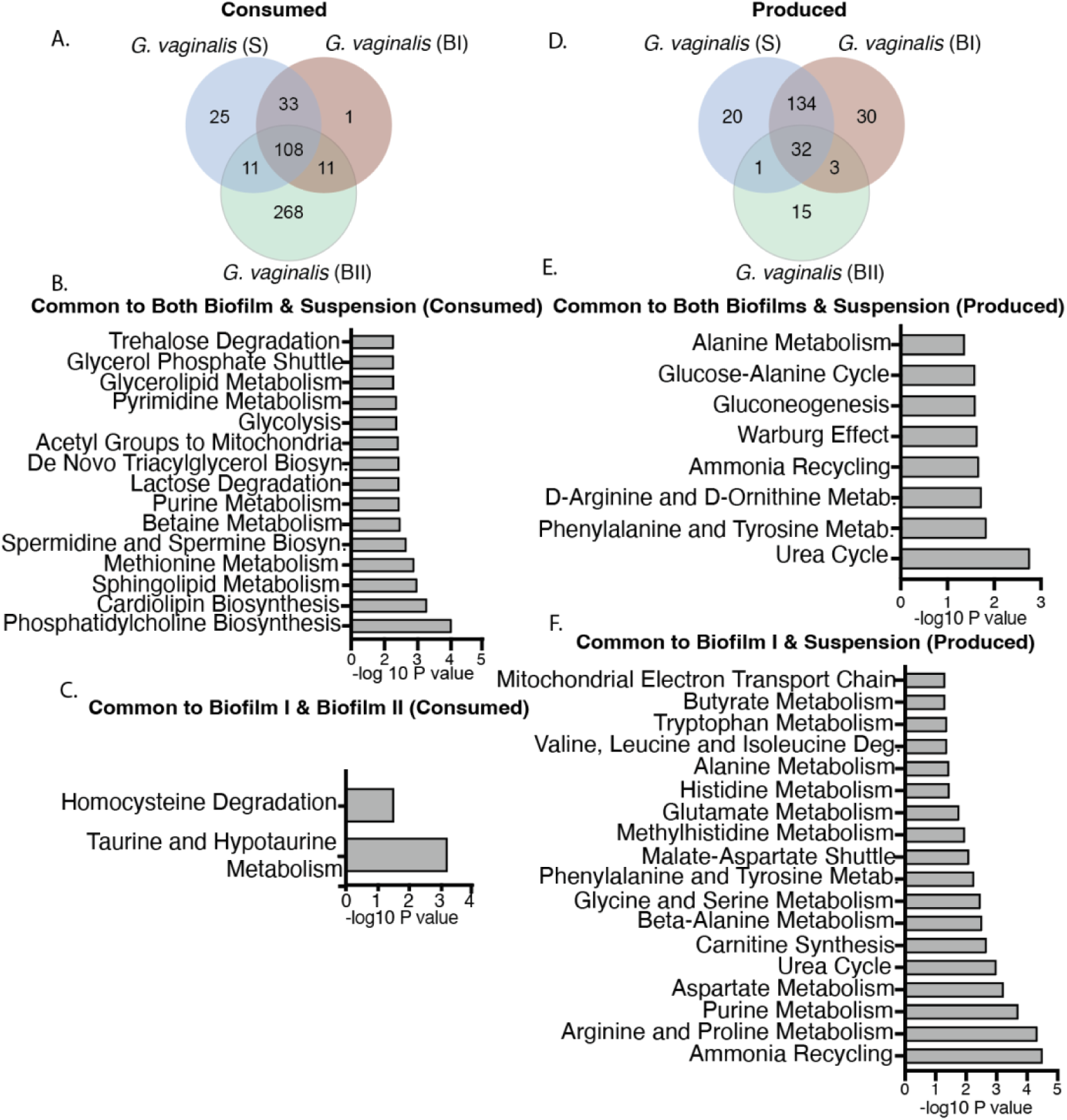
Shared metabolites across culture types of *G. vaginalis* indicate more similarity between suspension and biofilm type I and distinction between the two biofilm types. Intra-strain comparison of metabolite distribution and enrichment pathways corresponding to culture conditions of *G. vaginalis*. (A) Venn diagram showing distribution of consumed metabolites among suspension and biofilm cultures, (B) Common significant pathways for all biofilm and suspension consumed metabolites, (C) Pathways significant for both biofilm type I and biofilm type II based on consumed metabolites, (D) Venn diagram showing distribution of produced metabolites among suspension and biofilm cultures,(E) Common significant pathways for biofilm and suspension produced metabolites, (F) Common significant pathways for suspension and biofilm I produced metabolites. Included pathways with p value ≤ 0.059 and -log10 p value was used to plot the graphs. S: Suspension, BI: Biofilm type I, BII: Biofilm type II, Metab.: metabolism.

In terms of commonly produced metabolites, suspension cultures and biofilm type I shared 134 byproducts, while suspension cultures and biofilm type II shared only one. This indicates that biofilm type I is more similar to suspension culture in its metabolic profile compared to biofilm type II (Fig 2D). Also, only three commonly produced metabolites were shared between the two biofilm types, and these did not correspond to any specific significant pathway. This underscore distinct metabolic profiles that differentiate the two biofilm types. Among the enriched pathways associated with the produced metabolites, both suspension and type I biofilm cultures were linked to 18 common metabolic pathways, with ammonia recycling identified as the most significant (Fig 2F). However, with the inclusion of biofilm type II, urea cycle emerged as most predominant common pathway (Fig 2E). This suggests that biofilm type II is less efficient in conserving and reusing nitrogen compared to suspension cultures and biofilm type I. Pathways like D-ornithine metabolism, warburg effect and gluconeogenesis were only noticed when there was inclusion of biofilm type II along with suspension and type I biofilm (Fig 2E), (Supplementary table 1).

### Metabolites produced exclusively by suspension or biofilm cultures of *L. crispatus* belong to significantly different metabolic pathways

We discovered that *L. crispatus* suspension and biofilm cultures commonly consumed 142 metabolites and produced 104 metabolites, respectively (Fig 3A & 3E). The metabolites commonly consumed by both cultures were associated with six metabolic pathways, with methionine metabolism emerging as the most significantly abundant (Fig 3B). Metabolites exclusively consumed by suspension cultures also predominantly belonged to the methionine metabolism pathway, closely followed by spermidine and spermine biosynthesis (Fig 3C). In contrast, the metabolic profile of biofilm-exclusive consumed metabolites revealed a distinct set of pathways, most notably phenylacetate metabolism, followed by histidine metabolism, purine metabolism, and glutamate metabolism (Fig 3D).

**Fig 3.**
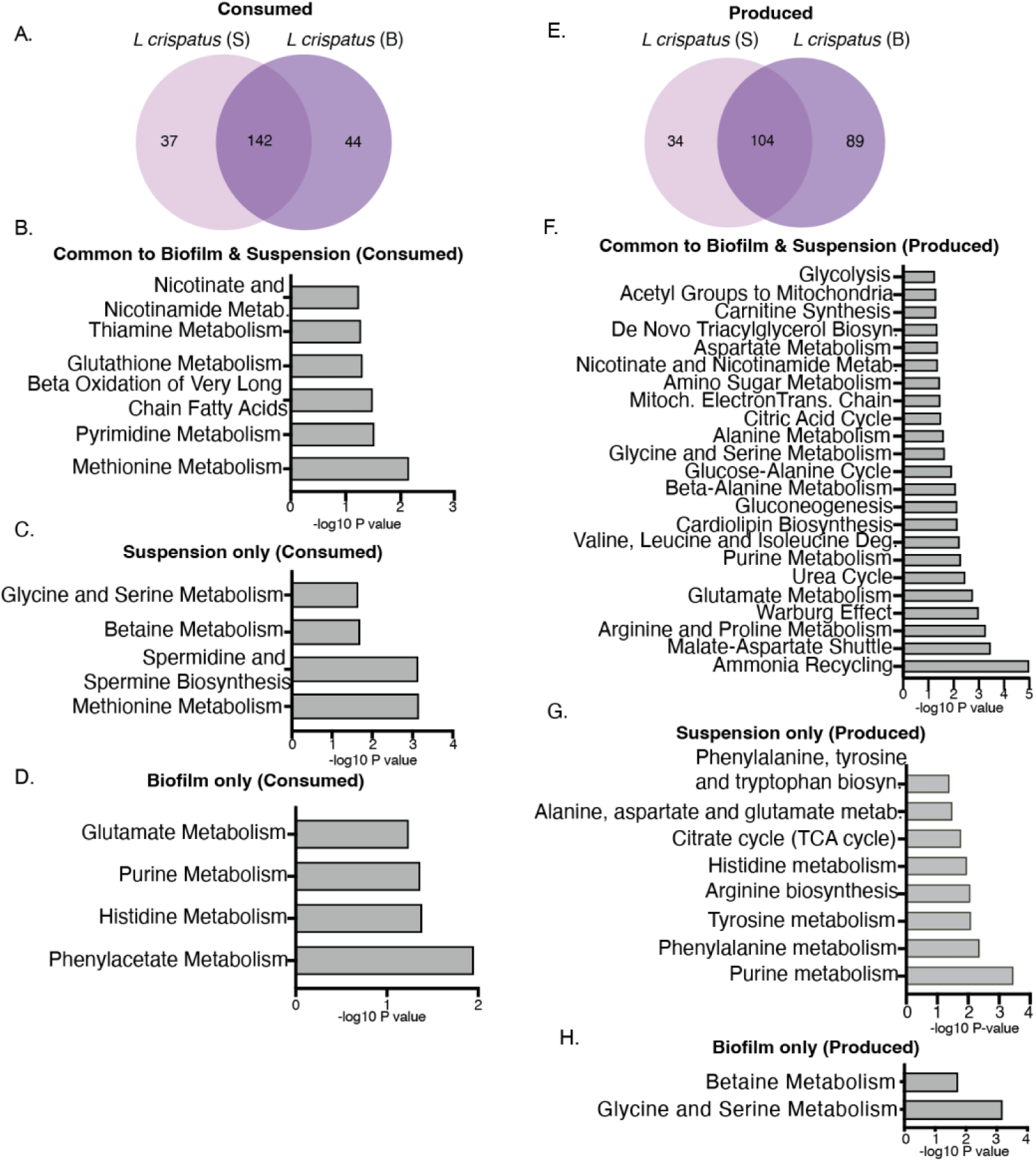
Suspension and biofilm populations of *L. crispatus* exhibit distinct metabolic profiles and have significant differences in key biochemical pathways. Intra-strain comparison of metabolite distribution and enrichment pathways corresponding to culture conditions of *L. crispatus*.(A) Venn diagram showing distribution of consumed metabolites among suspension and biofilm cultures, (B) significant pathways for biofilm and suspension shared consumed metabolites, (C) key metabolic pathways uniquely associated with suspension consumed metabolites, (D) key metabolic pathways uniquely associated with biofilm consumed metabolites, (E) Venn diagram showing distribution of produced metabolites among suspension and biofilm cultures, (F) significant pathways for biofilm and suspension shared produced metabolites, (G) key metabolic pathways uniquely associated with suspension produced metabolites, (H) key metabolic pathways uniquely associated with biofilm produced metabolites. Included pathways with p value ≤ 0.059 and -log10 p value was used to plot the graphs. S: Suspension, B: Biofilm, Metab.: metabolism, Deg.: Degradation, Biosyn.: Biosynthesis.

Metabolites commonly produced by both cultures were significantly associated with 22 metabolic pathways, with ammonia recycling emerging as the most prominent. This set of pathways was notably distinct from those observed in either culture individually (Fig 3F). The 34 metabolites exclusive to suspension culture highlighted purine metabolism as the most significant, followed by phenylalanine and tyrosine metabolism, arginine biosynthesis, histidine metabolism, and the TCA cycle (Fig 3G). In contrast, the biofilm culture of *L. crispatus* diverged from its suspension counterpart, primarily in glycine and serine metabolism and betaine metabolism pathways (Fig 3H), (Supplementary table 1).

### Commonly consumed metabolites in suspension or biofilm cultures of *L. crispatus, L. iners*, and *G. vaginalis* are predominantly associated with similar metabolic pathways, unlike produced metabolites

Our metabolomic analysis revealed 45 metabolites commonly consumed in suspension cultures of *L. crispatus, L. iners,* and *G. vaginalis* and 64 metabolites in biofilm cultures of *L. crispatus*, and both types of *G. vaginalis* biofilms (Fig 4A & B). Pathway enrichment analysis of these shared “consumed” metabolites in both culture types display methionine metabolism, phosphatidylcholine biosynthesis, purine metabolism, betaine metabolism, and mitochondrial acetyl group transfer as common pathways followed (Fig 4E & F). Suspension cultures differ from biofilm only by sphingolipid and methylhistidine metabolism pathways (Fig 4E). Glutathione metabolism emerged as the sole exclusive pathway enriched in biofilm cultures (Fig 4F). Conversely, the “produced” metabolites shared among suspension cultures were associated with distinct pathways compared to those produced by biofilm cultures (Fig 4G & H). Metabolites exclusively consumed by *L. iners* predominantly belonged to amino acid pathways, including glycine, serine, glutathione, and alanine metabolism, coupled with ammonia recycling processes. (Supplementary Fig 1a).

**Fig 4:**
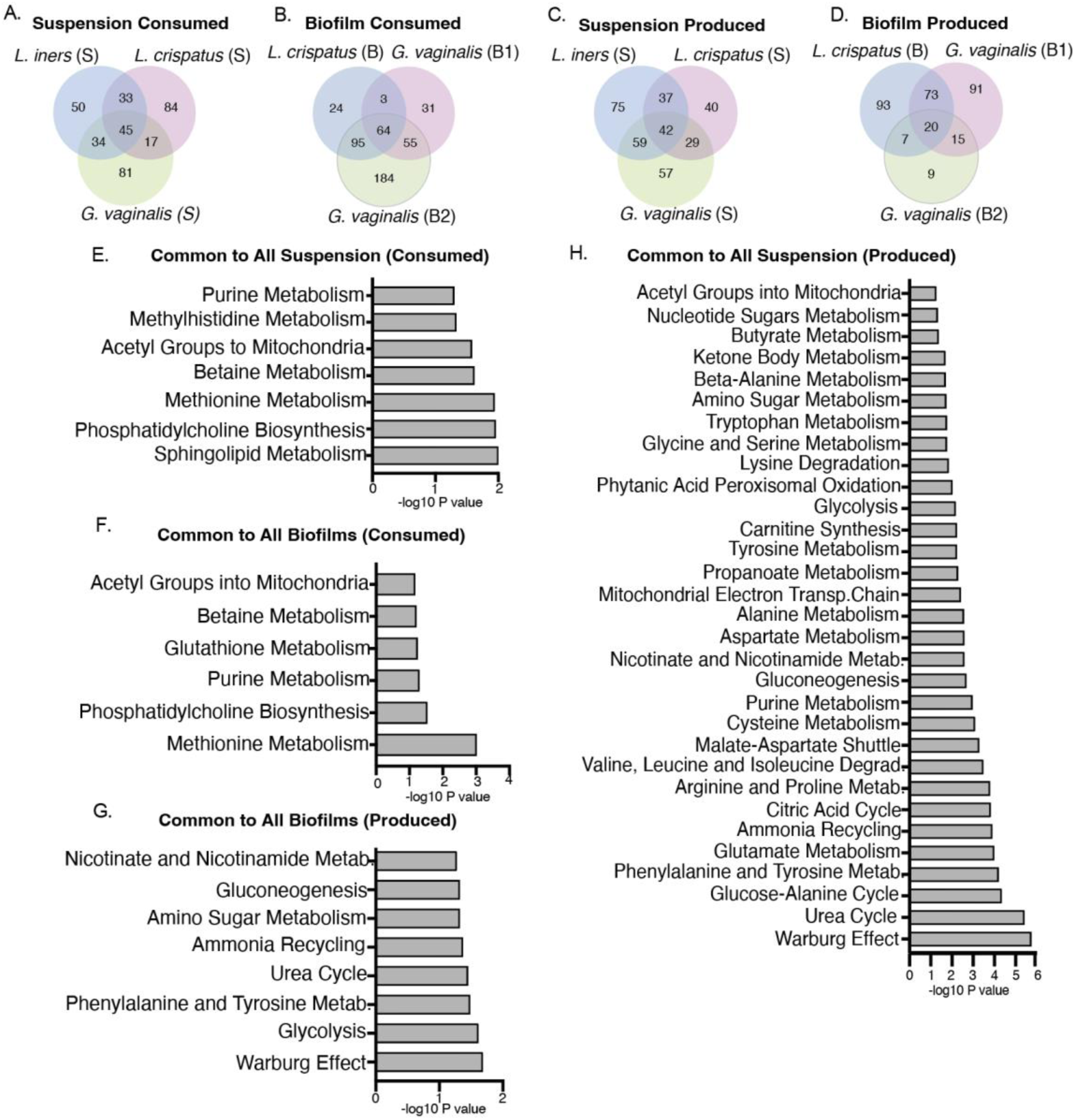
Biofilm produced metabolites belonged to more distinct pathways compared to suspension produced metabolites unlike consumed metabolites, that shows relatively more similar pathways between the two culture types Inter-microbial comparison of metabolite enrichment pathways of *L. crispatus, G. vaginalis*, *L. iners* suspension cultures and *L. crispatus* and *G. vaginalis* biofilm cultures (A-D) Venn diagrams showing distribution of metabolites common or exclusive for suspension and biofilm cultures, (E) significant pathways for all suspension consumed shared metabolites, (F) significant pathways for all biofilm consumed shared metabolites (G) significant pathways for all biofilm produced shared metabolites (H) significant pathways for all suspension produced shared metabolites. Included pathways with p value ≤ 0.059 and -log10 p value was used to plot the graphs. S: Suspension, B: Biofilm, B1: Biofilm type I, B2: Biofilm type II, metab.: Metabolism, Degrad.: Degradation.

Pathway enrichment analysis of the 42 suspension-produced common metabolites (Fig 4C) and 20 biofilm-produced metabolites (Fig 4D) revealed 31 and 8 enriched pathways followed respectively (Fig 4G & H). Suspension cultures followed 23 unique pathways than biofilm cultures (Fig 4G & H). This distribution highlights the metabolic divergence between suspension and biofilm growth modes with respect to produced metabolites unlike consumed metabolites that show more similarity. *L. iners* produces 75 unique metabolites largely associated with eight metabolic pathways, with homocysteine degradation being the most prominent, followed by taurine and hypotaurine metabolism (Supplementary Fig 1b), (Supplementary table 1).

### Certain metabolically essential host compounds were differentially produced or consumed by planktonic and/or biofilm cultures of key vaginal microbiota

Ophthalmate, accumulation of which is a biomarker of oxidative stress, was found to be significantly consumed by *L. crispatus* suspension and biofilm cultures only. Orotate, a precursor for pyrimidine biosynthesis was consumed by both suspension and biofilm cultures of *L. crispatus* and *G. vaginalis* biofilm type II. This compound was seen to be minimally produced by *L. iners* and *G. vaginalis* suspension and *G. vaginalis* biofilm type I (Fig 5A & B). Ribose was significantly produced by suspension culture of both the Lactobacillus spp. in this study but neither produced nor consumed by *G. vaginalis* (Fig 5A). Thymidine, crucial for DNA synthesis and repair, was exclusively utilized by *L. crispatus* biofilm (Fig 5B). Taurine, a sulfonic acid with critical functions in the central nervous system, exhibited contrasting metabolic patterns among vaginal microbiota. It was synthesized by *L. crispatus* biofilms while being significantly catabolized by *G. vaginalis* type II biofilms. (Fig. 5B).

**Fig 5.**
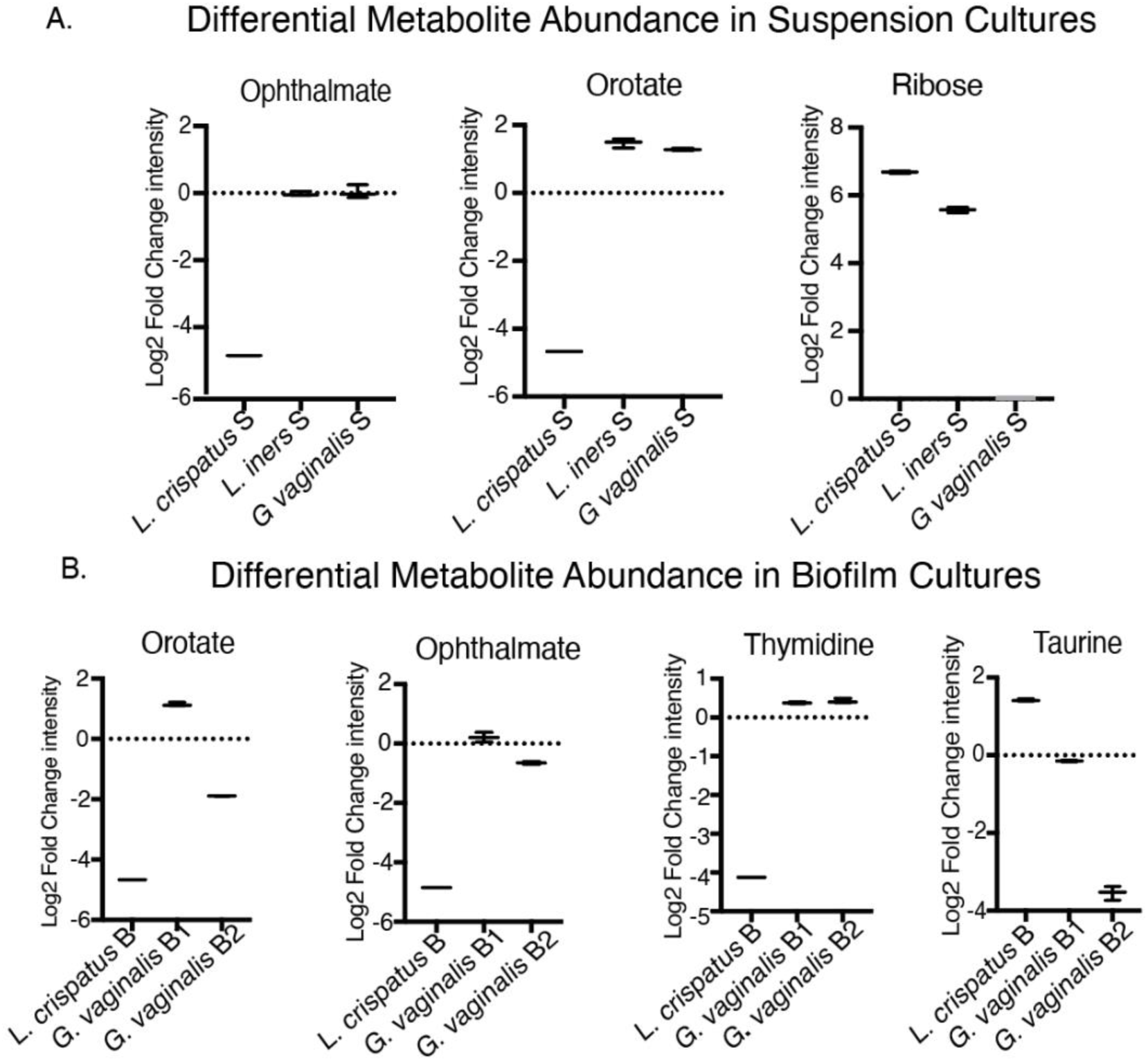
Differential abundance of representative key physiological metabolites among *L. crispatus, G. vaginalis,* and *L. iners* Cultures (A) Box plots of selected metabolites in suspension culture showing differential abundance between microbes, (B) Box plots of selected metabolites in biofilm culture showing differential abundance between microbes. Biological triplicate values were used to plot the graph. Log 2 fold change < 0: Consumed and > 0: Produced. S: Suspension, B: Biofilm, B1: Biofilm type I, B2: Biofilm type II.

## Discussions

In recent years, metabolomics has emerged as a potent tool, complementing other ‘omic’ studies and delving into the extensive metabolic diversity observed in prokaryotes ^12–14^. While some studies propose metabolomics as a more advanced and effective strategy for identifying BV, there is currently no study providing a comprehensive comparison of metabolite profiles among pure cultures of microbes significant to the vaginal environment ^44^. Additionally, to the best of our knowledge, there has been no research conducted to compare metabolic differences between planktonic and biofilm culture conditions of these microbes.

This study reveals contrasting metabolic patterns between planktonic and biofilm cultures of vaginal microbes, consistent with research indicating metabolic diversity among microbial cells within a biofilm matrix. Such heterogeneity arises from gradients of oxygen, pH, and nutrients within the biofilm, which are absent in planktonic cultures ^45^. The difference in metabolites produced by *G. vaginalis* biofilms with different media explains the metabolic shift in biofilm depending on nutrient availability, previously reported as a regulatory factor in biofilm development ^46^. The ability of *G. vaginalis* to thrive and proliferate with various available nutrients highlights its adaptable and resilient nature in biofilm formation.

In this study, both *L. crispatus* and *L. iners* were prominent producers of glutamine, a compound known to influence gut glutathione production, lymphocyte proliferation, and the secretion of certain cytokines ^47, 48^. Furthermore, these Lactobacillus *sp*. produced NAD^+^ with a notably higher level in *L. crispatus* biofilm cultures. Although limited research directly connects NAD^+^ levels in the vaginal microbiome to BV or fertility in women, studies suggest that certain gut bacteria can synthesize NAD^+^. This synthesis can influence the composition and function of the gut microbiota, affecting intestinal barrier function and immune modulation ^49, 50^. We hypothesize that these principles governing the gut microbiome ecosystem may also be applicable to the vaginal microenvironment, offering a compelling avenue for future scientific exploration and research. Key possibilities include: (1) Boosting NAD+ production within the vaginal microbiome may offer a novel strategy for maintaining balance and preventing bacterial vaginosis. (2) NAD^+^ might play a role in the communication between vaginal bacteria and host cells, potentially influencing fertility. We noticed gluconate production exclusively in *L. crispatus,* while both types of *G. vaginalis* biofilm consumed it. This aligns with previous reports indicating that pathogenic bacteria utilize and require gluconate for virulence and antibiotic resistance ^51, 52^. Furthermore, *L. crispatus*, was found to produce kynurenine, a signature compound in a healthy vaginal environment with higher levels in biofilm as compared to suspension culture ^53^. Substantial consumption of kynurenine by *G. vaginalis* in both suspension and biofilm forms, as well as by *L. iners* in suspension cultures was observed. Unlike *L. crispatus*, *L. iners* has been considered unfit in maintaining a healthy vaginal environment because for its lack of D-lactate production leading to poor pH maintenance, promoting pathogen adherence, lack of hydrogen peroxide production and pore forming protein, inerolysin production ^54^. In this study, *L. iners* were identified as the only significant producers of histamine that plays significant roles in reproductive physiology, including ovulation, embryo implantation, placental barrier regulation, lactation, and uterine contractions. Additionally, histamine dysregulation is implicated in pregnancy complications like pre-eclampsia and preterm birth ^55, 56, 57^. This suggests, although not the ideal microbe in the vaginal environment, it has potential involvement in modulating women’s reproductive health through its metabolites.

Metabolomic pathway analysis of consumed metabolites from *G. vaginalis* planktonic and biofilm cultures identified phosphatidylcholine (ChoP) biosynthesis as a key shared metabolic pathway. ChoP has been implicated in bacterial virulence and pathogenicity through its role in modulating host immune responses ^58^. Lecithinase, a virulence factor of *G. vaginalis* hydrolyzes ChoP, leading to reproductive cell and tissue damage ^59^. These findings highlight the complex interplay between *G. vaginalis* metabolism, virulence factors, and host interactions in the context of reproductive tract infections. Besides, we found that *G. vaginalis* consumes metabolites involved in cardiolipin biosynthesis pathway. This again, draws parallels to the intestinal pathogen *Shigella flexneri,* suggesting that cardiolipin might play a similar role in enhancing *G. vaginalis* virulence, possibly by supporting membrane integrity, protein localization, or other virulence-associated functions^60^. Studies have shown the role of cardiolipin in regulating bacterial respiratory chain and shape in *P. aeruginosa* and suggested targeting cardiolipin microdomains may be of great interest for developing new antibacterial therapies^61^. Our metabolomic analysis revealed contrasting patterns in taurine metabolism between *G. vaginalis* and *L. crispatus*. Both biofilm types of *G. vaginalis* consumed metabolites associated with the taurine metabolism pathway, whereas *L. crispatus* biofilms exhibited notable production of taurine. The role of taurine in vaginal health remains understudied, despite its significant functions in other physiological systems. This β-amino acid is known for its diverse biological roles, including neuromodulation in the central nervous system, regulation of renal function, antioxidant properties, and involvement in cellular metabolic and physiological processes^62,63^. This study suggests that taurine metabolism could be an important factor in the complex interplay between vaginal microbiota and host health.

We noticed striking differences in the metabolite profiles of the two types of *G. vaginalis* biofilms, with only two common metabolic pathways identified based on consumed metabolites. This observation highlights metabolic plasticity as an important adaptive strategy, allowing *G. vaginalis* to thrive in diverse nutrient environments. Such metabolic adaptability may contribute to the persistence and virulence of *G. vaginalis* in different host microenvironments. Understanding these metabolic adaptations could provide insights into *G. vaginalis* biofilm formation mechanisms.

This study represents a pioneering effort towards elucidating the specific growth requirements of these microbes that are significant to women’s health. Moreover, our data revealed a differential metabolite profile associated with critical metabolic pathways in biofilms of significant vaginal microbes compared to their planktonic cultures. Specifically, we identified metabolites exclusively required for the growth of *L. crispatus*, which could be leveraged to enhance the growth and stability of this beneficial microbe, thereby promoting a healthy vaginal environment. The metabolic pathways for the utilization of metabolites consumed exclusively by *G. vaginalis*, present potential targets for future therapeutic studies against BV. A deeper investigation will be pivotal to explore the therapeutic potential of these microbial metabolites for managing BV during pregnancy, where antibiotics are often avoided.

## Methods

### Bacterial culture

The microbial strains utilized in this study were *Lactobacillus crispatus* DSM20584, *Gardnerella vaginalis* ATCC 14018, and *Lactobacillus iners* AB107 (ATCC 55195), all procured from the American Type Culture Collection (ATCC). *L. crispatus* was cultivated in de Man, Rogosa and Sharpe (MRS) medium for both planktonic and biofilm cultures. *G. vaginalis* planktonic cultures were propagated in NYCIII medium, while biofilm formation was induced using two distinct substrates: NYCIII and a modified Brain Heart Infusion (sBHI) medium supplemented with 2% (w/v) gelatin, 0.5% (w/v) yeast extract, and 0.1% (w/v) soluble starch. *L. iners* planktonic cultures were grown in Brain Heart Infusion (BHI) medium. All cultures were incubated at 37°C in a 5% CO_2_, with planktonic cultures subjected to 24-hour agitation and biofilm cultures maintained under static conditions for 48 hours.

### Preparation of cell-free culture supernatant

Post incubation, cultures were centrifuged at 13000 x g for 10mins and then sterile filtered using 0.2μm filter. Cell-free supernatants were then subjected to non-targeted metabolomics in the Metabolon, NC, USA facility. Triplicates were used for each sample type.

### Non-targeted metabolomics

The comprehensive metabolomic profiling was conducted externally by Metabolon, Inc. Briefly, the sample preparation process utilized the automated MicroLab STAR® system from Hamilton Company. Recovery standards were introduced before initiating the extraction for quality control and the resulting extract was separated into five fractions. Organic solvent removal was performed using a TurboVap® (Zymark). To ensure data quality and reliability, various control types were analyzed in parallel with the experimental samples. UPLC-MS/MS was performed using a Waters ACQUITY UPLC platform interfaced with a Thermo Scientific Q-Exactive mass spectrometer, incorporating a HESI-II source and an Orbitrap analyzer with 35,000 mass resolution capability. Four distinct analytical methods were utilized as described by Metabolon. Metabolon’s proprietary hardware and software which relies on a robust library of authenticated standards for compound identification was used for the process of data extraction, compound identification, and quality control. Metabolite quantification involved area-under-the-curve analysis of peaks. Data normalization was achieved by adjusting run-day block medians to 1.00 and scaling each data point accordingly.

### Generation of heatmap

The intensity values of each metabolite for all the microbes and culture conditions were normalized with the respective blank media used for culturing them.

log2 Fold change (FC)= log2 (Mean_B_/ Mean_A_) (A is the blank media intensity value).

Log2 values were calculated and used to plot heatmaps using R studio. To mitigate computational constraints and enhance visualization clarity, a threshold-based data reduction approach was implemented. Specifically, only metabolites exhibiting a log2 fold change magnitude of ≥ 4 (i.e., log2 FC ≥ 4 or ≤ -4) were selected for inclusion in the heatmap generation process, thereby focusing on metabolites with substantial differential abundance.

### Intra and inter-microbial comparison of metabolites between suspension and biofilm cultures

Comparative metabolomic analysis was performed using Venn diagrams generated via the web-based tool at bioinformatics.psb.ugent.be to visualize the distribution of metabolites across *L. crispatus*, *G. vaginalis and L. iners* suspension and biofilm cultures (Only suspension culture for *L. iners*).

Metabolites unique to each intersection were subjected to metabolite set enrichment analysis (MSEA) using Metabo Analyst that used hypergeometric test to elucidate their respective metabolomic pathways. Pathway significance was visualized using bar diagrams created in GraphPad PRISM 10.0, with the y-axis representing -log10(p-value) for pathways meeting the significance threshold (p ≤ 0.059) (Supplementary table 1). Box plots were constructed to depict the log2 fold change in intensity of statistically significant metabolites (p value≤ 0.1), providing a quantitative representation of metabolite abundance variations across microbes and culture conditions (data submitted to Metabolomics Workbench).

### Data availability

The raw metabolomics data has been deposited in the Metabolomics Workbench under project number PR001854 and can be accessed via http://dx.doi.org/10.21228/M82M8R.

## Conclusion

This study represents a pioneering effort towards elucidating the specific growth requirements of these microbes that are significant to women’s health. Moreover, our data revealed a differential metabolite profile associated with critical metabolic pathways in biofilms of significant vaginal microbes compared to their planktonic cultures. Specifically, we identified metabolites exclusively required for the growth of *L. crispatus*, which could be leveraged to enhance the growth and stability of this beneficial microbe, thereby promoting a healthy vaginal environment. The metabolic pathways for the utilization of metabolites consumed exclusively by *G. vaginalis*, present potential targets for future therapeutic studies against BV.

## Acknowledgements

This work is supported by startup funding from the Weldon School of Biomedical Engineering, Purdue University and Department of Pathology at Case Western Reserve University.

## Author contributions

SJ & DKB conceived the study; SJ conducted experiments; SJ & DL analyzed the output data; SJ drafted the original manuscript, LNG & DKB reviewed and edited the draft. All authors have read and approved the final manuscript.

## Additional information

Supplementary files are submitted separately.

---

**Fig S1:**
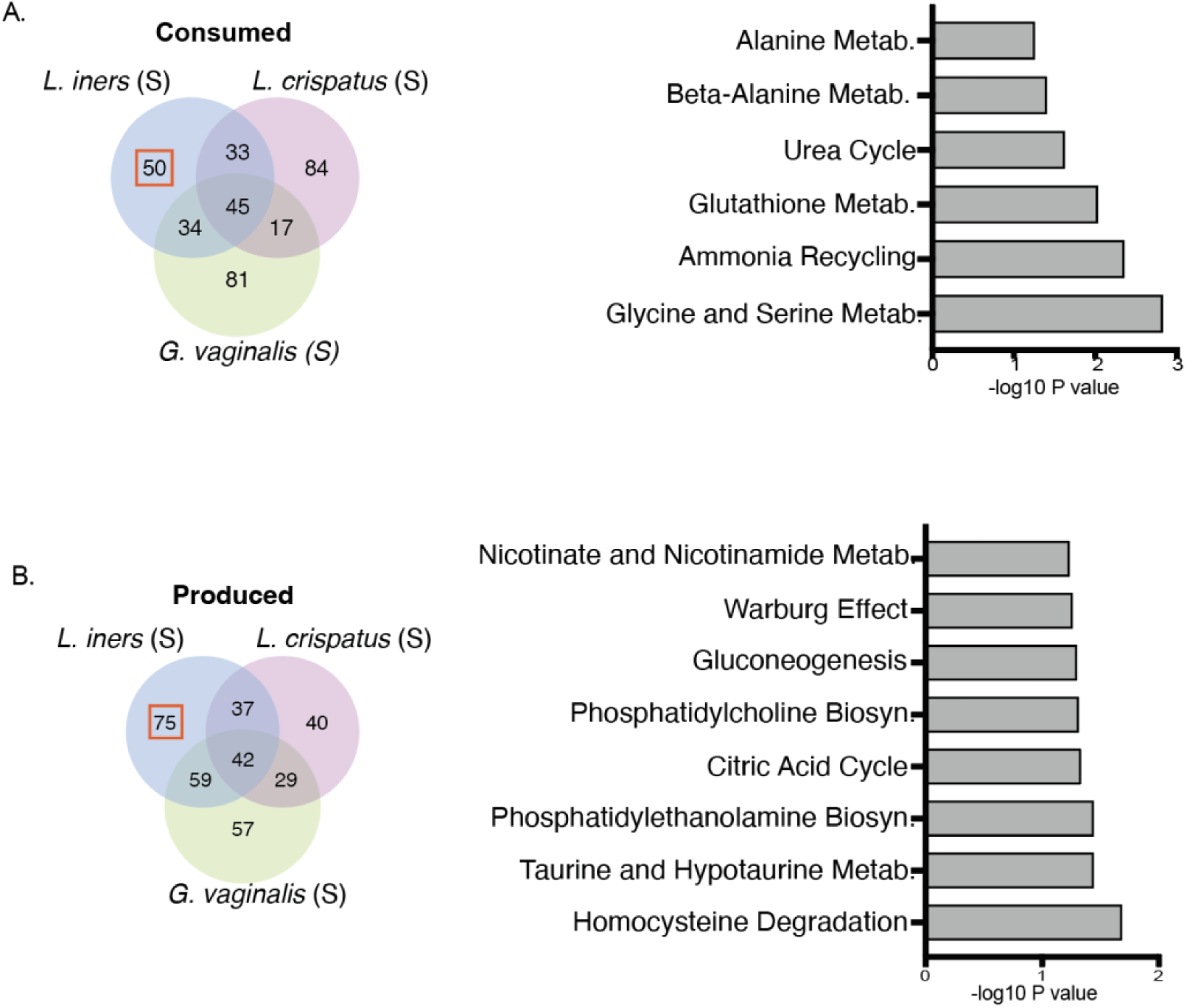
Suspension cultures of *L. iners* demonstrate distinctive metabolite profiles in terms of production and consumption, differing from *L. crispatus* and *G.vaginalis* cultures, with these metabolites linked to important metabolic pathways. (A) Venn diagram showing distribution of consumed metabolites common or exclusive for suspension cultures of *L. iners, L. crispatus* and *G. vaginalis* (left), significant pathways for metabolites consumed exclusively by *L. iners* (right). (B) Venn diagram showing distribution of produced metabolites common or exclusive for suspension cultures of *L. iners, L. crispatus* and *G. vaginalis* (left), significant pathways for metabolites produced exclusively by *L. iners* (right). Biological triplicate values were used to plot the graph. Included pathways with p value ≤ 0.059 and -log10 p value was used to plot the graphs. S: Suspension, Metab.: Metabolism, Biosyn.: Biosynthesis.

