## Supplementary Figure 1 for "Comparative Metabolomic Analysis of Vaginal Microbiota in Planktonic and Biofilm States Unveils Species-Specific Metabolic Signatures": Supplementary Fig 1.pdf

**Fig S1: Suspension cultures of *L. iners* demonstrate distinctive metabolite profiles in terms of production and consumption, differing from *L. crispatus* and *G. vaginalis* cultures, with these metabolites linked to important metabolic pathways.**

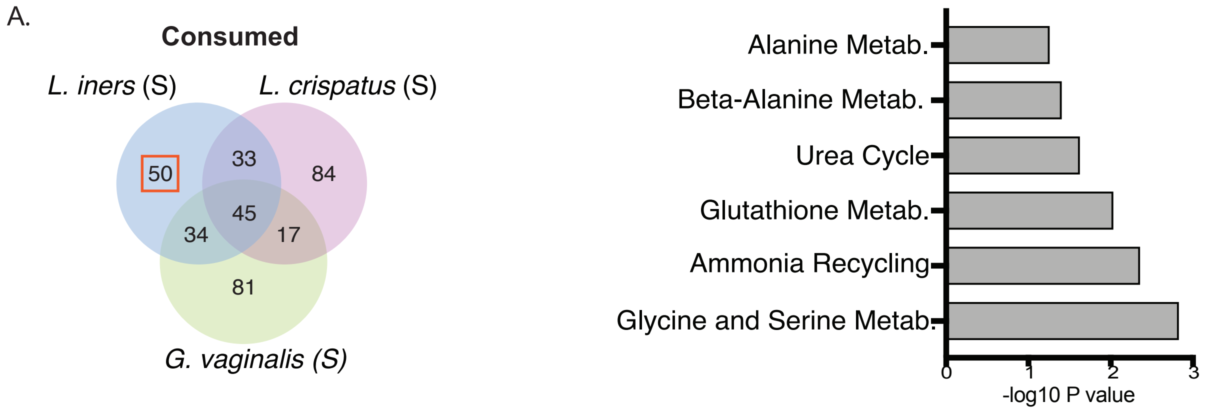


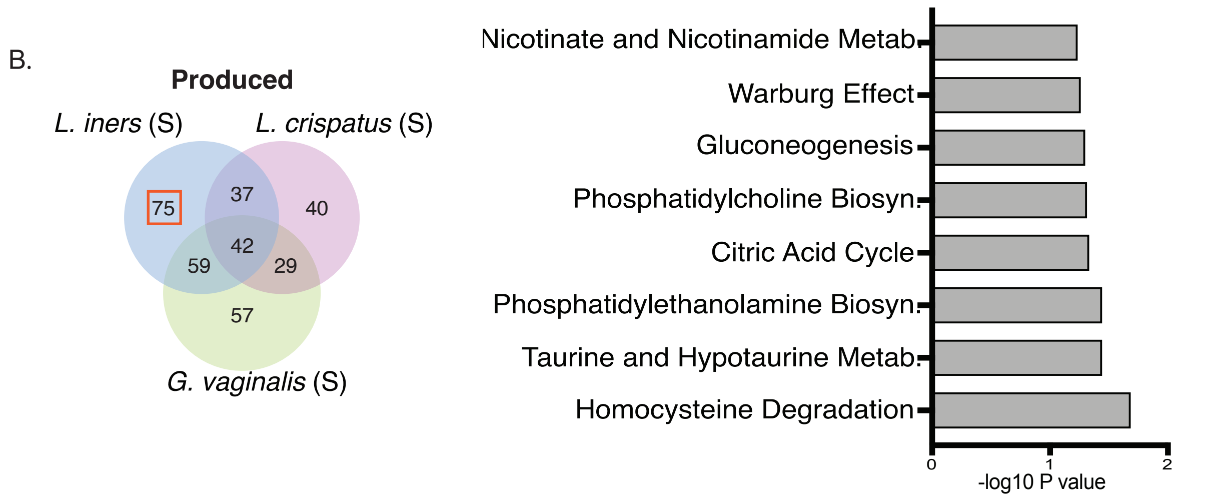


(A) Venn diagram showing distribution of consumed metabolites common or exclusive for suspension cultures of *L. iners, L. crispatus* and *G. vaginalis* ( left), significant pathways for metabolites consumed exclusively by *L. iners* (right). (B) Venn diagram showing distribution of produced metabolites common or exclusive for suspension cultures of *L. iners, L. crispatus* and *G. vaginalis* ( left), significant pathways for metabolites produced exclusively by *L. iners* (right).
Biological triplicate values were used to plot the graph. Included pathways with p value ≤ 0.059 and -log10 p value was used to plot the graphs. S: Suspension, Metab.: Metabolism, Biosyn.: Biosynthesis
