## Supplementary Table 1 for "Comparative Metabolomic Analysis of Vaginal Microbiota in Planktonic and Biofilm States Unveils Species-Specific Metabolic Signatures": Supplementary table 1.pdf

| Abbreviations | Complete term |
| --- | --- |
| S | Suspension |
| B | Biofilm |
| B1 | G. vaginalis biofilm type 1 |
| B2 | G. vaginalis biofilm type 2 |

| Consumed by all microbe B | | | |
|---|---|---|---|
| **Metabolic pathway** | **Raw p** | **FDR** | **minus log 10 p** |
| Methionine Metabolism | 0.000988 | 0.0969 | 3.005243055 |
| Phosphatidylcholine Biosynthesis | 0.0298 | 1 | 1.525783736 |
| Purine Metabolism | 0.0515 | 1 | 1.288192771 |
| Glutathione Metabolism | 0.058 | 1 | 1.236572006 |
| Betaine Metabolism | 0.0634 | 1 | 1.197910742 |
| Transfer of Acetyl Groups into Mitochondria | 0.0689 | 1 | 1.161780778 |
| Methylhistidine Metabolism | 0.0776 | 1 | 1.110138279 |
| Nicotinate and Nicotinamide Metabolism | 0.152 | 1 | 0.818156412 |
| Fatty Acid Biosynthesis | 0.152 | 1 | 0.818156412 |
| Phenylacetate Metabolism | 0.167 | 1 | 0.777283529 |
| Homocysteine Degradation | 0.167 | 1 | 0.777283529 |
| Lactose Degradation | 0.167 | 1 | 0.777283529 |
| Galactose Metabolism | 0.174 | 1 | 0.759450752 |
| Sphingolipid Metabolism | 0.188 | 1 | 0.725842151 |
| Trehalose Degradation | 0.2 | 1 | 0.698970004 |
| Histidine Metabolism | 0.203 | 1 | 0.692503962 |
| Taurine and Hypotaurine Metabolism | 0.216 | 1 | 0.665546249 |
| Glucose-Alanine Cycle | 0.232 | 1 | 0.634512015 |
| Beta Oxidation of Very Long Chain Fatty Acids | 0.292 | 1 | 0.534617149 |
| Spermidine and Spermine Biosynthesis | 0.307 | 1 | 0.512861625 |
| Warburg Effect | 0.316 | 1 | 0.500312917 |
| Butyrate Metabolism | 0.321 | 1 | 0.493494968 |
| Lactose Synthesis | 0.321 | 1 | 0.493494968 |
| Glycine and Serine Metabolism | 0.332 | 1 | 0.478861916 |
| Tryptophan Metabolism | 0.332 | 1 | 0.478861916 |
| Catecholamine Biosynthesis | 0.334 | 1 | 0.476253533 |
| Ubiquinone Biosynthesis | 0.334 | 1 | 0.476253533 |
| Pantothenate and CoA Biosynthesis | 0.348 | 1 | 0.458420756 |
| Carnitine Synthesis | 0.361 | 1 | 0.442492798 |
| Glycolysis | 0.374 | 1 | 0.427128398 |
| Estrone Metabolism | 0.387 | 1 | 0.412289035 |
| Cysteine Metabolism | 0.412 | 1 | 0.385102784 |
| Selenoamino Acid Metabolism | 0.424 | 1 | 0.372634143 |
| Mitochondrial Beta-Oxidation of Short Chain Saturated Fatty Aci | 0.424 | 1 | 0.372634143 |
| Phospholipid Biosynthesis | 0.447 | 1 | 0.349692477 |
| Starch and Sucrose Metabolism | 0.47 | 1 | 0.327902142 |
| Fructose and Mannose Degradation | 0.47 | 1 | 0.327902142 |
| Citric Acid Cycle | 0.481 | 1 | 0.317854924 |
| Amino Sugar Metabolism | 0.492 | 1 | 0.308034897 |
| Gluconeogenesis | 0.492 | 1 | 0.308034897 |
| Glutamate Metabolism | 0.629 | 1 | 0.201349355 |
| Arginine and Proline Metabolism | 0.659 | 1 | 0.181114585 |

| | | | |
|---|---|---|---|
| Pyrimidine Metabolism | 0.694 | 1 | 0.15864053 |
| Tyrosine Metabolism | 0.768 | 1 | 0.11463878 |

| Produced by L. iners S only | | | |
|---|---|---|---|
| **Metabolic pathway** | **Raw p** | **FDR** | **minus log 10 p** |
| Homocysteine Degradation | 0.0208 | 0.667 | 1.681936665 |
| Taurine and Hypotaurine Metabolism | 0.0364 | 0.667 | 1.438898616 |
| Phosphatidylethanolamine Biosynthesis | 0.0364 | 0.667 | 1.438898616 |
| Citric Acid Cycle | 0.0467 | 0.667 | 1.330683119 |
| Phosphatidylcholine Biosynthesis | 0.0487 | 0.667 | 1.312471039 |
| Gluconeogenesis | 0.0505 | 0.667 | 1.296708622 |
| Warburg Effect | 0.0551 | 0.667 | 1.258848401 |
| Nicotinate and Nicotinamide Metabolism | 0.0585 | 0.667 | 1.232844134 |
| Glycine and Serine Metabolism | 0.0613 | 0.667 | 1.212539525 |
| Vitamin B6 Metabolism | 0.0845 | 0.755 | 1.073143291 |
| Methionine Metabolism | 0.0911 | 0.755 | 1.040481623 |
| Glutathione Metabolism | 0.0924 | 0.755 | 1.034328029 |
| Betaine Metabolism | 0.101 | 0.758 | 0.995678626 |
| Glycolysis | 0.117 | 0.81 | 0.931814138 |
| Glutamate Metabolism | 0.124 | 0.81 | 0.906578315 |
| Arginine and Proline Metabolism | 0.148 | 0.906 | 0.829738285 |
| Urea Cycle | 0.162 | 0.935 | 0.790484985 |
| Biotin Metabolism | 0.19 | 1 | 0.721246399 |
| Thiamine Metabolism | 0.211 | 1 | 0.675717545 |
| Fatty Acid Biosynthesis | 0.229 | 1 | 0.640164518 |
| Ketone Body Metabolism | 0.291 | 1 | 0.536107011 |
| Histidine Metabolism | 0.298 | 1 | 0.525783736 |
| Pyruvate Metabolism | 0.347 | 1 | 0.459670525 |
| Alanine Metabolism | 0.363 | 1 | 0.440093375 |
| Spermidine and Spermine Biosynthesis | 0.38 | 1 | 0.420216403 |
| Catecholamine Biosynthesis | 0.412 | 1 | 0.385102784 |
| Riboflavin Metabolism | 0.412 | 1 | 0.385102784 |
| Threonine and 2-Oxobutanoate Degradation | 0.412 | 1 | 0.385102784 |
| Pantothenate and CoA Biosynthesis | 0.428 | 1 | 0.368556231 |
| Transfer of Acetyl Groups into Mitochondria | 0.443 | 1 | 0.353596274 |
| Glycerolipid Metabolism | 0.486 | 1 | 0.313363731 |
| Cysteine Metabolism | 0.5 | 1 | 0.301029996 |
| Phospholipid Biosynthesis | 0.539 | 1 | 0.268411235 |
| Ammonia Recycling | 0.563 | 1 | 0.249491605 |
| Starch and Sucrose Metabolism | 0.563 | 1 | 0.249491605 |
| Amino Sugar Metabolism | 0.586 | 1 | 0.232102384 |
| Beta-Alanine Metabolism | 0.597 | 1 | 0.224025669 |
| Aspartate Metabolism | 0.608 | 1 | 0.216096421 |
| Sphingolipid Metabolism | 0.658 | 1 | 0.181774106 |
| Propanoate Metabolism | 0.676 | 1 | 0.170053304 |
| Valine, Leucine and Isoleucine Degradation | 0.798 | 1 | 0.097997109 |
| Tryptophan Metabolism | 0.798 | 1 | 0.097997109 |

| | | | |
|---|---|---|---|
| Tyrosine Metabolism | 0.852 | 1 | 0.069560405 |
| Purine Metabolism | 0.864 | 1 | 0.063486258 |

| Consumed by L. iners S only | | | |
|---|---|---|---|
| **Metabolic pathways** | **Raw p** | **FDR** | **minus log 10 p** |
| Glycine and Serine Metabolism | 0.00149 | 0.146 | 2.826813732 |
| Ammonia Recycling | 0.00441 | 0.216 | 2.355561411 |
| Glutathione Metabolism | 0.00934 | 0.305 | 2.029653124 |
| Urea Cycle | 0.0238 | 0.583 | 1.623423043 |
| Beta-Alanine Metabolism | 0.0397 | 0.779 | 1.401209493 |
| Alanine Metabolism | 0.0556 | 0.908 | 1.254925208 |
| Purine Metabolism | 0.0798 | 1 | 1.097997109 |
| Methylhistidine Metabolism | 0.0888 | 1 | 1.051587034 |
| Glutamate Metabolism | 0.0932 | 1 | 1.030584088 |
| Glycerolipid Metabolism | 0.11 | 1 | 0.958607315 |
| Arginine and Proline Metabolism | 0.112 | 1 | 0.950781977 |
| Selenoamino Acid Metabolism | 0.125 | 1 | 0.903089987 |
| Tryptophan Metabolism | 0.149 | 1 | 0.826813732 |
| Homocysteine Degradation | 0.189 | 1 | 0.723538196 |
| Aspartate Metabolism | 0.19 | 1 | 0.721246399 |
| Malate-Aspartate Shuttle | 0.208 | 1 | 0.681936665 |
| Galactose Metabolism | 0.216 | 1 | 0.665546249 |
| Taurine and Hypotaurine Metabolism | 0.244 | 1 | 0.612610174 |
| Phosphatidylethanolamine Biosynthesis | 0.244 | 1 | 0.612610174 |
| Methionine Metabolism | 0.25 | 1 | 0.602059991 |
| Glucose-Alanine Cycle | 0.262 | 1 | 0.581698709 |
| Spermidine and Spermine Biosynthesis | 0.344 | 1 | 0.463441557 |
| Vitamin B6 Metabolism | 0.359 | 1 | 0.444905551 |
| Threonine and 2-Oxobutanoate Degradatior | 0.374 | 1 | 0.427128398 |
| Pantothenate and CoA Biosynthesis | 0.389 | 1 | 0.410050399 |
| Carnitine Synthesis | 0.403 | 1 | 0.394694954 |
| Inositol Metabolism | 0.507 | 1 | 0.294992041 |
| Starch and Sucrose Metabolism | 0.519 | 1 | 0.284832642 |
| Fructose and Mannose Degradation | 0.519 | 1 | 0.284832642 |
| Gluconeogenesis | 0.541 | 1 | 0.266802735 |
| Retinol Metabolism | 0.563 | 1 | 0.249491605 |
| Porphyrin Metabolism | 0.612 | 1 | 0.213248578 |
| Sphingolipid Metabolism | 0.612 | 1 | 0.213248578 |
| Histidine Metabolism | 0.631 | 1 | 0.199970641 |
| Pyruvate Metabolism | 0.673 | 1 | 0.171984936 |
| Pyrimidine Metabolism | 0.744 | 1 | 0.128427064 |
| Warburg Effect | 0.744 | 1 | 0.128427064 |
| Bile Acid Biosynthesis | 0.79 | 1 | 0.102372909 |
| Tyrosine Metabolism | 0.815 | 1 | 0.088842391 |

| Consumed by all microbes S | | | |
|---|---|---|---|
| **Metabolic Pathways** | **Raw p** | **FDR** | **minus log 10 p** |
| Sphingolipid Metabolism | 0.0101 | 0.378 | 1.995678626 |
| Phosphatidylcholine Biosynthesis | 0.0111 | 0.378 | 1.954677021 |
| Methionine Metabolism | 0.0116 | 0.378 | 1.935542011 |
| Betaine Metabolism | 0.0243 | 0.521 | 1.614393726 |
| Transfer of Acetyl Groups into Mitochondri | 0.0266 | 0.521 | 1.575118363 |
| Methylhistidine Metabolism | 0.0471 | 0.709 | 1.326979093 |
| Purine Metabolism | 0.0507 | 0.709 | 1.294992041 |
| Nicotinate and Nicotinamide Metabolism | 0.0628 | 0.77 | 1.202040356 |
| Galactose Metabolism | 0.0728 | 0.792 | 1.137868621 |
| Lactose Degradation | 0.103 | 1 | 0.987162775 |
| Trehalose Degradation | 0.125 | 1 | 0.903089987 |
| Warburg Effect | 0.146 | 1 | 0.835647144 |
| Glucose-Alanine Cycle | 0.146 | 1 | 0.835647144 |
| Spermidine and Spermine Biosynthesis | 0.196 | 1 | 0.707743929 |
| Lactose Synthesis | 0.206 | 1 | 0.68613278 |
| Catecholamine Biosynthesis | 0.216 | 1 | 0.665546249 |
| Ubiquinone Biosynthesis | 0.216 | 1 | 0.665546249 |
| Sulfate/Sulfite Metabolism | 0.235 | 1 | 0.628932138 |
| Carnitine Synthesis | 0.235 | 1 | 0.628932138 |
| Glycolysis | 0.244 | 1 | 0.612610174 |
| Estrone Metabolism | 0.254 | 1 | 0.595166283 |
| Selenoamino Acid Metabolism | 0.281 | 1 | 0.55129368 |
| Phospholipid Biosynthesis | 0.298 | 1 | 0.525783736 |
| Starch and Sucrose Metabolism | 0.316 | 1 | 0.500312917 |
| Fructose and Mannose Degradation | 0.316 | 1 | 0.500312917 |
| Citric Acid Cycle | 0.324 | 1 | 0.48945499 |
| Amino Sugar Metabolism | 0.332 | 1 | 0.478861916 |
| Androgen and Estrogen Metabolism | 0.332 | 1 | 0.478861916 |
| Gluconeogenesis | 0.332 | 1 | 0.478861916 |
| Histidine Metabolism | 0.404 | 1 | 0.393618635 |
| Arginine and Proline Metabolism | 0.474 | 1 | 0.324221658 |
| Pyrimidine Metabolism | 0.507 | 1 | 0.294992041 |
| Glycine and Serine Metabolism | 0.519 | 1 | 0.284832642 |
| Tryptophan Metabolism | 0.519 | 1 | 0.284832642 |
| Tyrosine Metabolism | 0.583 | 1 | 0.234331445 |

| Produced by all microbes B | | | |
|---|---|---|---|
| **Metabolic pathways** | **Raw p** | **FDR** | **minus log 10 P** |
| Warburg Effect | 0.0208 | 0.655 | 1.681936665 |
| Glycolysis | 0.0245 | 0.655 | 1.610833916 |
| Phenylalanine and Tyrosine Metabolism | 0.0331 | 0.655 | 1.480172006 |
| Urea Cycle | 0.0355 | 0.655 | 1.449771647 |
| Ammonia Recycling | 0.0428 | 0.655 | 1.368556231 |
| Amino Sugar Metabolism | 0.0481 | 0.655 | 1.317854924 |
| Gluconeogenesis | 0.0481 | 0.655 | 1.317854924 |
| Nicotinate and Nicotinamide Metabolism | 0.0535 | 0.655 | 1.271646218 |
| Galactose Metabolism | 0.0621 | 0.676 | 1.2069084 |
| Glutamate Metabolism | 0.0938 | 0.846 | 1.027797162 |
| Phenylacetate Metabolism | 0.0949 | 0.846 | 1.022733788 |
| Pyruvaldehyde Degradation | 0.105 | 0.857 | 0.978810701 |
| D-Arginine and D-Ornithine Metabolism | 0.115 | 0.866 | 0.93930216 |
| Valine, Leucine and Isoleucine Degradatio | 0.133 | 0.878 | 0.876148359 |
| Glucose-Alanine Cycle | 0.134 | 0.878 | 0.872895202 |
| Alanine Metabolism | 0.172 | 1 | 0.764471553 |
| Purine Metabolism | 0.188 | 1 | 0.725842151 |
| Nucleotide Sugars Metabolism | 0.2 | 1 | 0.698970004 |
| Transfer of Acetyl Groups into Mitochondri | 0.218 | 1 | 0.661543506 |
| Inositol Phosphate Metabolism | 0.235 | 1 | 0.628932138 |
| Cysteine Metabolism | 0.252 | 1 | 0.598599459 |
| Pentose Phosphate Pathway | 0.277 | 1 | 0.557520231 |
| Inositol Metabolism | 0.285 | 1 | 0.54515514 |
| Starch and Sucrose Metabolism | 0.294 | 1 | 0.53165267 |
| Fructose and Mannose Degradation | 0.294 | 1 | 0.53165267 |
| Citric Acid Cycle | 0.302 | 1 | 0.519993057 |
| Aspartate Metabolism | 0.325 | 1 | 0.488116639 |
| Pyruvate Metabolism | 0.412 | 1 | 0.385102784 |
| Pyrimidine Metabolism | 0.477 | 1 | 0.321481621 |
| Glycine and Serine Metabolism | 0.489 | 1 | 0.310691141 |
| Tyrosine Metabolism | 0.551 | 1 | 0.258848401 |

| Produced by all microbes S | | | |
|---|---|---|---|
| **Metabolic pathways** | **Raw p** | **FDR** | **minus log 10 P** |
| Warburg Effect | 1.88E-06 | 0.000185 | 5.725842151 |
| Urea Cycle | 4.01E-06 | 0.000197 | 5.396855627 |
| Glucose-Alanine Cycle | 4.73E-05 | 0.00155 | 4.325138859 |
| Phenylalanine and Tyrosine Metabolism | 6.52E-05 | 0.0016 | 4.185752404 |
| Glutamate Metabolism | 0.000106 | 0.00206 | 3.974694135 |
| Ammonia Recycling | 0.000131 | 0.00206 | 3.882728704 |
| Citric Acid Cycle | 0.000154 | 0.00206 | 3.812479279 |
| Arginine and Proline Metabolism | 0.000168 | 0.00206 | 3.774690718 |
| Valine, Leucine and Isoleucine Degradation | 0.000345 | 0.00376 | 3.462180905 |
| Malate-Aspartate Shuttle | 0.000541 | 0.0053 | 3.266802735 |
| Cysteine Metabolism | 0.000854 | 0.00761 | 3.068542129 |
| Purine Metabolism | 0.00112 | 0.00916 | 2.950781977 |
| Gluconeogenesis | 0.00216 | 0.0163 | 2.665546249 |
| Nicotinate and Nicotinamide Metabolism | 0.0027 | 0.0173 | 2.568636236 |
| Aspartate Metabolism | 0.0027 | 0.0173 | 2.568636236 |
| Alanine Metabolism | 0.00283 | 0.0173 | 2.548213564 |
| Mitochondrial Electron Transport Chain | 0.00395 | 0.0227 | 2.403402904 |
| Propanoate Metabolism | 0.00533 | 0.029 | 2.273272791 |
| Tyrosine Metabolism | 0.00604 | 0.0297 | 2.218963061 |
| Carnitine Synthesis | 0.00606 | 0.0297 | 2.217527376 |
| Glycolysis | 0.00689 | 0.0322 | 2.161780778 |
| Phytanic Acid Peroxisomal Oxidation | 0.00978 | 0.0436 | 2.009661145 |
| Lysine Degradation | 0.0146 | 0.0622 | 1.835647144 |
| Glycine and Serine Metabolism | 0.0178 | 0.0699 | 1.749579998 |
| Tryptophan Metabolism | 0.0178 | 0.0699 | 1.749579998 |
| Amino Sugar Metabolism | 0.019 | 0.0715 | 1.721246399 |
| Beta-Alanine Metabolism | 0.0206 | 0.074 | 1.68613278 |
| Ketone Body Metabolism | 0.0212 | 0.074 | 1.673664139 |
| Butyrate Metabolism | 0.0435 | 0.147 | 1.361510743 |
| Nucleotide Sugars Metabolism | 0.0478 | 0.156 | 1.320572103 |
| Transfer of Acetyl Groups into Mitochondria | 0.0569 | 0.18 | 1.244887734 |
| Inositol Phosphate Metabolism | 0.0666 | 0.204 | 1.176525771 |
| Oxidation of Branched Chain Fatty Acids | 0.0768 | 0.228 | 1.11463878 |
| Folate Metabolism | 0.093 | 0.268 | 1.031517051 |
| Inositol Metabolism | 0.0986 | 0.276 | 1.006123085 |
| Starch and Sucrose Metabolism | 0.104 | 0.284 | 0.982966661 |
| Galactose Metabolism | 0.146 | 0.38 | 0.835647144 |
| Phenylacetate Metabolism | 0.151 | 0.38 | 0.821023053 |
| De Novo Triacylglycerol Biosynthesis | 0.151 | 0.38 | 0.821023053 |
| Pyruvaldehyde Degradation | 0.166 | 0.408 | 0.779891912 |
| Histidine Metabolism | 0.172 | 0.411 | 0.764471553 |
| Glycerol Phosphate Shuttle | 0.182 | 0.414 | 0.739928612 |

| | | | |
|---|---|---|---|
| Cardiolipin Biosynthesis | 0.182 | 0.414 | 0.739928612 |
| Pyruvate Metabolism | 0.205 | 0.457 | 0.688246139 |
| Thyroid hormone synthesis | 0.211 | 0.46 | 0.675717545 |
| Pyrimidine Metabolism | 0.273 | 0.582 | 0.563837353 |
| Ethanol Degradation | 0.294 | 0.601 | 0.53165267 |
| Catecholamine Biosynthesis | 0.307 | 0.601 | 0.512861625 |
| Glutathione Metabolism | 0.307 | 0.601 | 0.512861625 |
| Threonine and 2-Oxobutanoate Degradation | 0.307 | 0.601 | 0.512861625 |
| Betaine Metabolism | 0.319 | 0.613 | 0.496209317 |
| Caffeine Metabolism | 0.344 | 0.643 | 0.463441557 |
| Androstenedione Metabolism | 0.356 | 0.643 | 0.448550002 |
| Estrone Metabolism | 0.356 | 0.643 | 0.448550002 |
| Glycerolipid Metabolism | 0.368 | 0.643 | 0.434152181 |
| Plasmalogen Synthesis | 0.38 | 0.643 | 0.420216403 |
| Selenoamino Acid Metabolism | 0.391 | 0.643 | 0.407823243 |
| Mitochondrial Beta-Oxidation of Short Chain Saturated Fatty Acids | 0.391 | 0.643 | 0.407823243 |
| Mitochondrial Beta-Oxidation of Medium Chain Saturated Fatty Acid | 0.391 | 0.643 | 0.407823243 |
| Pterine Biosynthesis | 0.402 | 0.643 | 0.395773947 |
| Mitochondrial Beta-Oxidation of Long Chain Saturated Fatty Acids | 0.402 | 0.643 | 0.395773947 |
| Phospholipid Biosynthesis | 0.413 | 0.643 | 0.384049948 |
| Pentose Phosphate Pathway | 0.413 | 0.643 | 0.384049948 |
| Fructose and Mannose Degradation | 0.435 | 0.666 | 0.361510743 |
| Androgen and Estrogen Metabolism | 0.456 | 0.687 | 0.341035157 |
| Fatty Acid Elongation In Mitochondria | 0.476 | 0.696 | 0.322393047 |
| Retinol Metabolism | 0.476 | 0.696 | 0.322393047 |
| Porphyrin Metabolism | 0.523 | 0.753 | 0.281498311 |
| Methionine Metabolism | 0.54 | 0.758 | 0.26760624 |
| Fatty acid Metabolism | 0.549 | 0.758 | 0.260427656 |
| Steroidogenesis | 0.549 | 0.758 | 0.260427656 |
| Bile Acid Biosynthesis | 0.704 | 0.958 | 0.152427341 |
| Arachidonic Acid Metabolism | 0.715 | 0.96 | 0.145693958 |

| Consumed by G. vaginalis B1 & B2 | | | |
|---|---|---|---|
| **Metabolic pathways** | **Raw p** | **FDR** | **minus log 10 p** |
| Taurine and Hypotaurine Metabolism | 0.000779 | 0.0764 | 3.108462542 |
| Homocysteine Degradation | 0.0355 | 1 | 1.449771647 |
| Glutathione Metabolism | 0.0776 | 1 | 1.110138279 |
| Pantothenate and CoA Biosynthesis | 0.0814 | 1 | 1.089375595 |
| Cysteine Metabolism | 0.1 | 1 | 1 |
| Nicotinate and Nicotinamide Metabolism | 0.133 | 1 | 0.876148359 |
| Methionine Metabolism | 0.158 | 1 | 0.801342913 |
| Steroid Biosynthesis | 0.179 | 1 | 0.747146969 |
| Glutamate Metabolism | 0.179 | 1 | 0.747146969 |
| Glycine and Serine Metabolism | 0.216 | 1 | 0.665546249 |
| Tryptophan Metabolism | 0.216 | 1 | 0.665546249 |

| Consumed by S, B1 & B2 of G. vaginalis | | | |
|---|---|---|---|
| **Metabolic pathways** | **Raw p** | **FDR** | **minus log 10 p** |
| Phosphatidylcholine Biosynthesis | 0.000976 | 0.0957 | 3.010550182 |
| Cardiolipin Biosynthesis | 0.00532 | 0.261 | 2.274088368 |
| Sphingolipid Metabolism | 0.0105 | 0.317 | 1.978810701 |
| Methionine Metabolism | 0.0129 | 0.317 | 1.88941029 |
| Spermidine and Spermine Biosynthesis | 0.0222 | 0.35 | 1.653647026 |
| Betaine Metabolism | 0.0337 | 0.35 | 1.472370099 |
| Purine Metabolism | 0.0362 | 0.35 | 1.441291429 |
| Lactose Degradation | 0.0366 | 0.35 | 1.436518915 |
| De Novo Triacylglycerol Biosynthesis | 0.0366 | 0.35 | 1.436518915 |
| Transfer of Acetyl Groups into Mitochondria | 0.0381 | 0.35 | 1.419075024 |
| Glycolysis | 0.0428 | 0.35 | 1.368556231 |
| Pyrimidine Metabolism | 0.0435 | 0.35 | 1.361510743 |
| Glycerolipid Metabolism | 0.053 | 0.35 | 1.27572413 |
| Glycerol Phosphate Shuttle | 0.0535 | 0.35 | 1.271646218 |
| Trehalose Degradation | 0.0535 | 0.35 | 1.271646218 |
| Phosphatidylethanolamine Biosynthesis | 0.0628 | 0.385 | 1.202040356 |
| Starch and Sucrose Metabolism | 0.0899 | 0.518 | 1.046240308 |
| Gluconeogenesis | 0.104 | 0.547 | 0.982966661 |
| Phosphatidylinositol Phosphate Metabolism | 0.116 | 0.547 | 0.935542011 |
| Nicotinate and Nicotinamide Metabolism | 0.119 | 0.547 | 0.924453039 |
| Warburg Effect | 0.132 | 0.547 | 0.879426069 |
| Methylhistidine Metabolism | 0.133 | 0.547 | 0.876148359 |
| Mitochondrial Electron Transport Chain | 0.14 | 0.547 | 0.853871964 |
| Lactose Synthesis | 0.14 | 0.547 | 0.853871964 |
| Galactose Metabolism | 0.143 | 0.547 | 0.844663963 |
| Glycine and Serine Metabolism | 0.145 | 0.547 | 0.838631998 |
| Glutathione Metabolism | 0.152 | 0.552 | 0.818156412 |
| Sulfate/Sulfite Metabolism | 0.177 | 0.6 | 0.752026734 |
| Histidine Metabolism | 0.177 | 0.6 | 0.752026734 |
| Bile Acid Biosynthesis | 0.187 | 0.61 | 0.728158393 |
| Plasmalogen Synthesis | 0.229 | 0.725 | 0.640164518 |
| Selenoamino Acid Metabolism | 0.242 | 0.728 | 0.616184634 |
| Urea Cycle | 0.256 | 0.728 | 0.591760035 |
| Phospholipid Biosynthesis | 0.269 | 0.728 | 0.57024772 |
| Arginine and Proline Metabolism | 0.271 | 0.728 | 0.567030709 |
| Thiamine Metabolism | 0.275 | 0.728 | 0.560667306 |
| Phenylacetate Metabolism | 0.275 | 0.728 | 0.560667306 |
| Fructose and Mannose Degradation | 0.295 | 0.762 | 0.530177984 |
| Citric Acid Cycle | 0.309 | 0.776 | 0.510041521 |
| Amino Sugar Metabolism | 0.322 | 0.789 | 0.492144128 |
| Tryptophan Metabolism | 0.34 | 0.795 | 0.468521083 |
| Fatty Acid Biosynthesis | 0.348 | 0.795 | 0.458420756 |

| | | | |
|---|---|---|---|
| Taurine and Hypotaurine Metabolism | 0.349 | 0.795 | 0.457174573 |
| Glucose-Alanine Cycle | 0.372 | 0.828 | 0.42945706 |
| Propanoate Metabolism | 0.437 | 0.951 | 0.359518563 |
| Beta Oxidation of Very Long Chain Fatty Acids | 0.456 | 0.951 | 0.341035157 |
| Alanine Metabolism | 0.456 | 0.951 | 0.341035157 |
| Vitamin B6 Metabolism | 0.494 | 0.965 | 0.306273051 |
| Butyrate Metabolism | 0.494 | 0.965 | 0.306273051 |
| Catecholamine Biosynthesis | 0.512 | 0.965 | 0.290730039 |
| Ubiquinone Biosynthesis | 0.512 | 0.965 | 0.290730039 |
| Riboflavin Metabolism | 0.512 | 0.965 | 0.290730039 |
| Pantothenate and CoA Biosynthesis | 0.53 | 0.979 | 0.27572413 |
| Carnitine Synthesis | 0.546 | 0.992 | 0.262807357 |
| Inositol Phosphate Metabolism | 0.578 | 1 | 0.238072162 |
| Estrone Metabolism | 0.578 | 1 | 0.238072162 |
| Cysteine Metabolism | 0.608 | 1 | 0.216096421 |
| Mitochondrial Beta-Oxidation of Short Chain Saturated Fatty Aci | 0.622 | 1 | 0.206209615 |
| Pentose Phosphate Pathway | 0.649 | 1 | 0.187755303 |
| Folate Metabolism | 0.649 | 1 | 0.187755303 |
| Inositol Metabolism | 0.661 | 1 | 0.179798541 |
| Ammonia Recycling | 0.674 | 1 | 0.171340103 |
| Androgen and Estrogen Metabolism | 0.697 | 1 | 0.156767222 |
| Beta-Alanine Metabolism | 0.708 | 1 | 0.149966742 |
| Aspartate Metabolism | 0.718 | 1 | 0.143875556 |
| Pyruvate Metabolism | 0.819 | 1 | 0.086716098 |
| Steroid Biosynthesis | 0.826 | 1 | 0.083019953 |
| Glutamate Metabolism | 0.826 | 1 | 0.083019953 |
| Valine, Leucine and Isoleucine Degradation | 0.885 | 1 | 0.053056729 |
| Tyrosine Metabolism | 0.924 | 1 | 0.034328029 |

| Produced by S, B1 & B2 of G. vaginalis | | | |
|---|---|---|---|
| **Metabolic pathway** | **Raw p** | **FDR** | **minus log 10 P** |
| Urea Cycle | 0.00174 | 0.171 | 2.759450752 |
| Phenylalanine and Tyrosine Metabolism | 0.0147 | 0.362 | 1.832682665 |
| D-Arginine and D-Ornithine Metabolism | 0.0187 | 0.362 | 1.728158393 |
| Ammonia Recycling | 0.0214 | 0.362 | 1.669586227 |
| Warburg Effect | 0.023 | 0.362 | 1.638272164 |
| Gluconeogenesis | 0.0253 | 0.362 | 1.596879479 |
| Glucose-Alanine Cycle | 0.0259 | 0.362 | 1.586700236 |
| Alanine Metabolism | 0.043 | 0.527 | 1.366531544 |
| Glutamate Metabolism | 0.0663 | 0.722 | 1.178486472 |
| Glycolysis | 0.0745 | 0.731 | 1.127843727 |
| Cysteine Metabolism | 0.0924 | 0.758 | 1.034328029 |
| Pyrimidine Metabolism | 0.1 | 0.758 | 1 |
| Valine, Leucine and Isoleucine Degradation | 0.108 | 0.758 | 0.966576245 |
| Tryptophan Metabolism | 0.108 | 0.758 | 0.966576245 |
| Citric Acid Cycle | 0.131 | 0.781 | 0.882728704 |
| Amino Sugar Metabolism | 0.138 | 0.781 | 0.860120914 |
| Beta-Alanine Metabolism | 0.145 | 0.781 | 0.838631998 |
| Nicotinate and Nicotinamide Metabolism | 0.152 | 0.781 | 0.818156412 |
| Aspartate Metabolism | 0.152 | 0.781 | 0.818156412 |
| Phenylacetate Metabolism | 0.167 | 0.781 | 0.777283529 |
| Galactose Metabolism | 0.174 | 0.781 | 0.759450752 |
| Malate-Aspartate Shuttle | 0.183 | 0.781 | 0.73754891 |
| Pyruvaldehyde Degradation | 0.183 | 0.781 | 0.73754891 |
| Arginine and Proline Metabolism | 0.278 | 1 | 0.555955204 |
| Spermidine and Spermine Biosynthesis | 0.307 | 1 | 0.512861625 |
| Glycine and Serine Metabolism | 0.332 | 1 | 0.478861916 |
| Nucleotide Sugars Metabolism | 0.334 | 1 | 0.476253533 |
| Pantothenate and CoA Biosynthesis | 0.348 | 1 | 0.458420756 |
| Carnitine Synthesis | 0.361 | 1 | 0.442492798 |
| Transfer of Acetyl Groups into Mitochondria | 0.361 | 1 | 0.442492798 |
| Inositol Phosphate Metabolism | 0.387 | 1 | 0.412289035 |
| Oxidation of Branched Chain Fatty Acids | 0.412 | 1 | 0.385102784 |
| Phytanic Acid Peroxisomal Oxidation | 0.412 | 1 | 0.385102784 |
| Tyrosine Metabolism | 0.413 | 1 | 0.384049948 |
| Purine Metabolism | 0.435 | 1 | 0.361510743 |
| Pentose Phosphate Pathway | 0.447 | 1 | 0.349692477 |
| Inositol Metabolism | 0.459 | 1 | 0.338187314 |
| Lysine Degradation | 0.459 | 1 | 0.338187314 |
| Starch and Sucrose Metabolism | 0.47 | 1 | 0.327902142 |
| Fructose and Mannose Degradation | 0.47 | 1 | 0.327902142 |
| Propanoate Metabolism | 0.579 | 1 | 0.237321436 |
| Histidine Metabolism | 0.579 | 1 | 0.237321436 |

| | | | |
|---|---|---|---|
| Pyruvate Metabolism | 0.621 | 1 | 0.2069084 |

| Produced by S and B1 of G. vaginalis | | | |
|---|---|---|---|
| **Metabolic pathway** | **Raw p** | **FDR** | **minus log 10 p** |
| Ammonia Recycling | 3.19E-05 | 0.0023 | 4.496209317 |
| Arginine and Proline Metabolism | 4.69E-05 | 0.0023 | 4.328827157 |
| Purine Metabolism | 0.000197 | 0.00644 | 3.705533774 |
| Aspartate Metabolism | 0.000596 | 0.0146 | 3.22475374 |
| Urea Cycle | 0.00104 | 0.0205 | 2.982966661 |
| Carnitine Synthesis | 0.00217 | 0.0354 | 2.663540266 |
| Beta-Alanine Metabolism | 0.00304 | 0.0422 | 2.517126416 |
| Glycine and Serine Metabolism | 0.00344 | 0.0422 | 2.463441557 |
| Phenylalanine and Tyrosine Metabolism | 0.00561 | 0.0611 | 2.251037139 |
| Malate-Aspartate Shuttle | 0.00815 | 0.0798 | 2.088842391 |
| Methylhistidine Metabolism | 0.0112 | 0.0996 | 1.950781977 |
| Glutamate Metabolism | 0.0172 | 0.14 | 1.764471553 |
| Histidine Metabolism | 0.0357 | 0.259 | 1.447331784 |
| Alanine Metabolism | 0.037 | 0.259 | 1.431798276 |
| Valine, Leucine and Isoleucine Degradation | 0.0434 | 0.266 | 1.36251027 |
| Tryptophan Metabolism | 0.0434 | 0.266 | 1.36251027 |
| Butyrate Metabolism | 0.0496 | 0.27 | 1.304518324 |
| Mitochondrial Electron Transport Chain | 0.0496 | 0.27 | 1.304518324 |
| Pyrimidine Metabolism | 0.106 | 0.44 | 0.974694135 |
| Warburg Effect | 0.106 | 0.44 | 0.974694135 |
| Cysteine Metabolism | 0.107 | 0.44 | 0.970616222 |
| Ketone Body Metabolism | 0.112 | 0.44 | 0.950781977 |
| Glucose-Alanine Cycle | 0.112 | 0.44 | 0.950781977 |
| Propanoate Metabolism | 0.114 | 0.44 | 0.943095149 |
| Methionine Metabolism | 0.114 | 0.44 | 0.943095149 |
| Selenoamino Acid Metabolism | 0.117 | 0.44 | 0.931814138 |
| Lysine Degradation | 0.148 | 0.536 | 0.829738285 |
| Bile Acid Biosynthesis | 0.161 | 0.563 | 0.793174124 |
| Citric Acid Cycle | 0.17 | 0.574 | 0.769551079 |
| Tyrosine Metabolism | 0.2 | 0.638 | 0.698970004 |
| Nicotinate and Nicotinamide Metabolism | 0.205 | 0.638 | 0.688246139 |
| Ethanol Degradation | 0.208 | 0.638 | 0.681936665 |
| Glutathione Metabolism | 0.225 | 0.65 | 0.647817482 |
| Threonine and 2-Oxobutanoate Degradation | 0.225 | 0.65 | 0.647817482 |
| Biotin Metabolism | 0.309 | 0.767 | 0.510041521 |
| Glycerolipid Metabolism | 0.311 | 0.767 | 0.507239611 |
| Oxidation of Branched Chain Fatty Acids | 0.328 | 0.767 | 0.484126156 |
| Phytanic Acid Peroxisomal Oxidation | 0.328 | 0.767 | 0.484126156 |
| Thiamine Metabolism | 0.34 | 0.767 | 0.468521083 |
| Phenylacetate Metabolism | 0.34 | 0.767 | 0.468521083 |
| Homocysteine Degradation | 0.34 | 0.767 | 0.468521083 |
| De Novo Triacylglycerol Biosynthesis | 0.34 | 0.767 | 0.468521083 |

| | | | |
|---|---|---|---|
| Mitochondrial Beta-Oxidation of Short Chain Saturated Fatty Acids | 0.344 | 0.767 | 0.463441557 |
| Mitochondrial Beta-Oxidation of Medium Chain Saturated Fatty Aci | 0.344 | 0.767 | 0.463441557 |
| Pyruvate Metabolism | 0.354 | 0.77 | 0.450996738 |
| Mitochondrial Beta-Oxidation of Long Chain Saturated Fatty Acids | 0.361 | 0.77 | 0.442492798 |
| Phospholipid Biosynthesis | 0.378 | 0.771 | 0.4225082 |
| Folate Metabolism | 0.378 | 0.771 | 0.4225082 |
| Glycerol Phosphate Shuttle | 0.398 | 0.781 | 0.400116928 |
| Cardiolipin Biosynthesis | 0.398 | 0.781 | 0.400116928 |
| Phosphatidylethanolamine Biosynthesis | 0.426 | 0.818 | 0.370590401 |
| Gluconeogenesis | 0.442 | 0.834 | 0.354577731 |
| Thyroid hormone synthesis | 0.452 | 0.835 | 0.344861565 |
| Retinol Metabolism | 0.473 | 0.859 | 0.325138859 |
| Beta Oxidation of Very Long Chain Fatty Acids | 0.545 | 0.941 | 0.263603498 |
| Porphyrin Metabolism | 0.546 | 0.941 | 0.262807357 |
| Fatty acid Metabolism | 0.587 | 0.941 | 0.231361899 |
| Steroidogenesis | 0.587 | 0.941 | 0.231361899 |
| Nucleotide Sugars Metabolism | 0.605 | 0.941 | 0.218244625 |
| Catecholamine Biosynthesis | 0.605 | 0.941 | 0.218244625 |
| Riboflavin Metabolism | 0.605 | 0.941 | 0.218244625 |
| Pantothenate and CoA Biosynthesis | 0.623 | 0.941 | 0.205511953 |
| Betaine Metabolism | 0.623 | 0.941 | 0.205511953 |
| Sulfate/Sulfite Metabolism | 0.64 | 0.941 | 0.193820026 |
| Transfer of Acetyl Groups into Mitochondria | 0.64 | 0.941 | 0.193820026 |
| Caffeine Metabolism | 0.657 | 0.941 | 0.18243463 |
| Glycolysis | 0.657 | 0.941 | 0.18243463 |
| Inositol Phosphate Metabolism | 0.672 | 0.941 | 0.172630727 |
| Androstenedione Metabolism | 0.672 | 0.941 | 0.172630727 |
| Estrone Metabolism | 0.672 | 0.941 | 0.172630727 |
| Plasmalogen Synthesis | 0.702 | 0.969 | 0.153662888 |
| Pterine Biosynthesis | 0.729 | 0.986 | 0.137272472 |
| Pentose Phosphate Pathway | 0.741 | 0.986 | 0.130181792 |
| Inositol Metabolism | 0.753 | 0.986 | 0.123205024 |
| Starch and Sucrose Metabolism | 0.765 | 0.986 | 0.116338565 |
| Fructose and Mannose Degradation | 0.765 | 0.986 | 0.116338565 |
| Amino Sugar Metabolism | 0.786 | 0.988 | 0.104577454 |
| Androgen and Estrogen Metabolism | 0.786 | 0.988 | 0.104577454 |
| Fatty Acid Elongation In Mitochondria | 0.805 | 0.999 | 0.09420412 |
| Galactose Metabolism | 0.831 | 1 | 0.080398976 |
| Sphingolipid Metabolism | 0.847 | 1 | 0.07211659 |
| Arachidonic Acid Metabolism | 0.959 | 1 | 0.018181393 |

| Consumed by both S & B of L. crispatus | | | |
|---|---|---|---|
| **Metabolic pathways** | **Raw p** | **FDR** | **minus log 10 P** |
| Methionine Metabolism | 0.00719 | 0.704 | 2.14327111 |
| Pyrimidine Metabolism | 0.0306 | 0.786 | 1.514278574 |
| Beta Oxidation of Very Long Chain Fatty Acids | 0.0329 | 0.786 | 1.482804102 |
| Glutathione Metabolism | 0.0504 | 0.786 | 1.297569464 |
| Thiamine Metabolism | 0.0535 | 0.786 | 1.271646218 |
| Nicotinate and Nicotinamide Metabolism | 0.0584 | 0.786 | 1.233587153 |
| Carnitine Synthesis | 0.0642 | 0.786 | 1.192464972 |
| Transfer of Acetyl Groups into Mitochondria | 0.0642 | 0.786 | 1.192464972 |
| Purine Metabolism | 0.0855 | 0.926 | 1.068033885 |
| Mitochondrial Beta-Oxidation of Short Chain Saturated Fatty Acid | 0.105 | 0.926 | 0.978810701 |
| Phosphatidylcholine Biosynthesis | 0.118 | 0.926 | 0.928117993 |
| Citric Acid Cycle | 0.154 | 0.926 | 0.812479279 |
| Methylhistidine Metabolism | 0.161 | 0.926 | 0.793174124 |
| Alanine Metabolism | 0.163 | 0.926 | 0.787812396 |
| Spermidine and Spermine Biosynthesis | 0.179 | 0.926 | 0.747146969 |
| Arginine and Proline Metabolism | 0.179 | 0.926 | 0.747146969 |
| Fatty Acid Biosynthesis | 0.186 | 0.926 | 0.730487056 |
| Butyrate Metabolism | 0.194 | 0.926 | 0.71219827 |
| Riboflavin Metabolism | 0.21 | 0.926 | 0.677780705 |
| Galactose Metabolism | 0.22 | 0.926 | 0.657577319 |
| Warburg Effect | 0.224 | 0.926 | 0.649751982 |
| Pantothenate and CoA Biosynthesis | 0.227 | 0.926 | 0.643974143 |
| Betaine Metabolism | 0.227 | 0.926 | 0.643974143 |
| Glycine and Serine Metabolism | 0.243 | 0.926 | 0.614393726 |
| Histidine Metabolism | 0.267 | 0.926 | 0.573488739 |
| Biotin Metabolism | 0.297 | 0.926 | 0.527243551 |
| Cysteine Metabolism | 0.308 | 0.926 | 0.511449283 |
| Oxidation of Branched Chain Fatty Acids | 0.308 | 0.926 | 0.511449283 |
| Phenylalanine and Tyrosine Metabolism | 0.324 | 0.926 | 0.48945499 |
| Selenoamino Acid Metabolism | 0.324 | 0.926 | 0.48945499 |
| Phenylacetate Metabolism | 0.327 | 0.926 | 0.485452247 |
| Homocysteine Degradation | 0.327 | 0.926 | 0.485452247 |
| Lactose Degradation | 0.327 | 0.926 | 0.485452247 |
| Glutamate Metabolism | 0.34 | 0.926 | 0.468521083 |
| Urea Cycle | 0.34 | 0.926 | 0.468521083 |
| Mitochondrial Beta-Oxidation of Long Chain Saturated Fatty Acids | 0.34 | 0.926 | 0.468521083 |
| Pentose Phosphate Pathway | 0.356 | 0.944 | 0.448550002 |
| Trehalose Degradation | 0.384 | 0.951 | 0.415668776 |
| Ammonia Recycling | 0.388 | 0.951 | 0.411168274 |
| Starch and Sucrose Metabolism | 0.388 | 0.951 | 0.411168274 |
| Taurine and Hypotaurine Metabolism | 0.411 | 0.978 | 0.386158178 |
| Gluconeogenesis | 0.419 | 0.978 | 0.377785977 |

| | | | |
|---|---|---|---|
| Glucose-Alanine Cycle | 0.437 | 0.995 | 0.359518563 |
| Aspartate Metabolism | 0.449 | 1 | 0.347753659 |
| Tryptophan Metabolism | 0.471 | 1 | 0.326979093 |
| Sphingolipid Metabolism | 0.521 | 1 | 0.283162277 |
| Propanoate Metabolism | 0.548 | 1 | 0.261219442 |
| Fatty acid Metabolism | 0.561 | 1 | 0.251037139 |
| Vitamin B6 Metabolism | 0.569 | 1 | 0.244887734 |
| Lactose Synthesis | 0.569 | 1 | 0.244887734 |
| Ethanol Degradation | 0.569 | 1 | 0.244887734 |
| Catecholamine Biosynthesis | 0.588 | 1 | 0.230622674 |
| Ubiquinone Biosynthesis | 0.588 | 1 | 0.230622674 |
| Threonine and 2-Oxobutanoate Degradation | 0.588 | 1 | 0.230622674 |
| Pyruvate Metabolism | 0.611 | 1 | 0.21395879 |
| Glycolysis | 0.64 | 1 | 0.193820026 |
| Estrone Metabolism | 0.655 | 1 | 0.1837587 |
| Mitochondrial Beta-Oxidation of Medium Chain Saturated Fatty A[illegible] | 0.699 | 1 | 0.155522824 |
| Phospholipid Biosynthesis | 0.725 | 1 | 0.139661993 |
| Lysine Degradation | 0.737 | 1 | 0.132532512 |
| Fructose and Mannose Degradation | 0.749 | 1 | 0.125518182 |
| Amino Sugar Metabolism | 0.77 | 1 | 0.113509275 |
| Bile Acid Biosynthesis | 0.784 | 1 | 0.105683937 |
| Valine, Leucine and Isoleucine Degradation | 0.931 | 1 | 0.031050319 |
| Tyrosine Metabolism | 0.959 | 1 | 0.018181393 |

| Consumed only by L. crispatus S | | | |
|---|---|---|---|
| **Metabolic pathways** | **Raw p** | **FDR** | **minus log 10 p** |
| Methionine Metabolism | 0.000713 | 0.036 | 3.14691047 |
| Spermidine and Spermine Biosynthesis | 0.000736 | 0.036 | 3.133122186 |
| Betaine Metabolism | 0.0205 | 0.559 | 1.688246139 |
| Glycine and Serine Metabolism | 0.0228 | 0.559 | 1.642065153 |
| Homocysteine Degradation | 0.0949 | 1 | 1.022733788 |
| D-Arginine and D-Ornithine Metabolism | 0.115 | 1 | 0.93930216 |
| Taurine and Hypotaurine Metabolism | 0.125 | 1 | 0.903089987 |
| Phosphatidylethanolamine Biosynthesis | 0.125 | 1 | 0.903089987 |
| Phosphatidylcholine Biosynthesis | 0.144 | 1 | 0.841637508 |
| Sulfate/Sulfite Metabolism | 0.218 | 1 | 0.661543506 |
| Cysteine Metabolism | 0.252 | 1 | 0.598599459 |
| Phenylalanine and Tyrosine Metabolism | 0.261 | 1 | 0.583359493 |
| Urea Cycle | 0.269 | 1 | 0.57024772 |
| Phospholipid Biosynthesis | 0.277 | 1 | 0.557520231 |
| Ammonia Recycling | 0.294 | 1 | 0.53165267 |
| Androgen and Estrogen Metabolism | 0.309 | 1 | 0.510041521 |
| Nicotinate and Nicotinamide Metabolism | 0.325 | 1 | 0.488116639 |
| Aspartate Metabolism | 0.325 | 1 | 0.488116639 |
| Sphingolipid Metabolism | 0.363 | 1 | 0.440093375 |
| Arginine and Proline Metabolism | 0.445 | 1 | 0.351639989 |
| Tryptophan Metabolism | 0.489 | 1 | 0.310691141 |
| Tyrosine Metabolism | 0.551 | 1 | 0.258848401 |

| Consumed by L. crispatus B only | | | |
|---|---|---|---|
| **Metabolic pathways** | **Raw p** | **FDR** | **minus log 10 p** |
| Phenylacetate Metabolism | 0.0113 | 1 | 1.946921557 |
| Histidine Metabolism | 0.0416 | 1 | 1.380906669 |
| Purine Metabolism | 0.0435 | 1 | 1.361510743 |
| Glutamate Metabolism | 0.0583 | 1 | 1.234331445 |
| Methylhistidine Metabolism | 0.0738 | 1 | 1.131943638 |
| Glycerolipid Metabolism | 0.0789 | 1 | 1.102922997 |
| Phenylalanine and Tyrosine Metabolism | 0.0902 | 1 | 1.044793462 |
| Tryptophan Metabolism | 0.0959 | 1 | 1.018181393 |
| Urea Cycle | 0.0961 | 1 | 1.017276612 |
| Aspartate Metabolism | 0.14 | 1 | 0.853871964 |
| Biotin Metabolism | 0.142 | 1 | 0.847711656 |
| Thiamine Metabolism | 0.159 | 1 | 0.798602876 |
| Methionine Metabolism | 0.187 | 1 | 0.728158393 |
| Arginine and Proline Metabolism | 0.259 | 1 | 0.586700236 |
| Alanine Metabolism | 0.28 | 1 | 0.552841969 |
| Vitamin B6 Metabolism | 0.307 | 1 | 0.512861625 |
| Butyrate Metabolism | 0.307 | 1 | 0.512861625 |
| Mitochondrial Electron Transport Chain | 0.307 | 1 | 0.512861625 |
| Ethanol Degradation | 0.307 | 1 | 0.512861625 |
| Glycine and Serine Metabolism | 0.309 | 1 | 0.510041521 |
| Glutathione Metabolism | 0.321 | 1 | 0.493494968 |
| Riboflavin Metabolism | 0.321 | 1 | 0.493494968 |
| Pantothenate and CoA Biosynthesis | 0.334 | 1 | 0.476253533 |
| Carnitine Synthesis | 0.347 | 1 | 0.459670525 |
| Tyrosine Metabolism | 0.388 | 1 | 0.411168274 |
| Cysteine Metabolism | 0.396 | 1 | 0.402304814 |
| Selenoamino Acid Metabolism | 0.408 | 1 | 0.389339837 |
| Mitochondrial Beta-Oxidation of Short Chain Saturated Fatty Acids | 0.408 | 1 | 0.389339837 |
| Mitochondrial Beta-Oxidation of Medium Chain Saturated Fatty Ac | 0.408 | 1 | 0.389339837 |
| Mitochondrial Beta-Oxidation of Long Chain Saturated Fatty Acids | 0.419 | 1 | 0.377785977 |
| Pentose Phosphate Pathway | 0.431 | 1 | 0.36552273 |
| Lysine Degradation | 0.442 | 1 | 0.354577731 |
| Ammonia Recycling | 0.453 | 1 | 0.343901798 |
| Citric Acid Cycle | 0.463 | 1 | 0.334419009 |
| Beta-Alanine Metabolism | 0.484 | 1 | 0.315154638 |
| Nicotinate and Nicotinamide Metabolism | 0.494 | 1 | 0.306273051 |
| Retinol Metabolism | 0.494 | 1 | 0.306273051 |
| Galactose Metabolism | 0.524 | 1 | 0.280668713 |
| Propanoate Metabolism | 0.56 | 1 | 0.251811973 |
| Fatty acid Metabolism | 0.569 | 1 | 0.244887734 |
| Pyruvate Metabolism | 0.602 | 1 | 0.220403509 |
| Warburg Effect | 0.675 | 1 | 0.170696227 |

| | | | |
|---|---|---|---|
| Bile Acid Biosynthesis | 0.724 | 1 | 0.140261434 |
| Arachidonic Acid Metabolism | 0.735 | 1 | 0.133712661 |

| Produced by both S & B of L. crispatus | | | |
|---|---|---|---|
| **Metabolic pathways** | **Raw p** | **FDR** | **minus log 10 p** |
| Ammonia Recycling | 1.03E-05 | 0.00101 | 4.987162775 |
| Malate-Aspartate Shuttle | 0.000349 | 0.0171 | 3.457174573 |
| Arginine and Proline Metabolism | 0.000541 | 0.0177 | 3.266802735 |
| Warburg Effect | 0.00103 | 0.0252 | 2.987162775 |
| Glutamate Metabolism | 0.00177 | 0.0346 | 2.752026734 |
| Urea Cycle | 0.0035 | 0.0572 | 2.455931956 |
| Purine Metabolism | 0.00527 | 0.0717 | 2.278189385 |
| Valine, Leucine and Isoleucine Degradation | 0.00595 | 0.0717 | 2.225483034 |
| Cardiolipin Biosynthesis | 0.00725 | 0.0717 | 2.139661993 |
| Gluconeogenesis | 0.00732 | 0.0717 | 2.135488919 |
| Beta-Alanine Metabolism | 0.00834 | 0.0743 | 2.078833949 |
| Glucose-Alanine Cycle | 0.0119 | 0.0972 | 1.924453039 |
| Glycine and Serine Metabolism | 0.0229 | 0.173 | 1.640164518 |
| Alanine Metabolism | 0.0254 | 0.178 | 1.595166283 |
| Citric Acid Cycle | 0.0321 | 0.205 | 1.493494968 |
| Mitochondrial Electron Transport Chain | 0.0343 | 0.205 | 1.46470588 |
| Amino Sugar Metabolism | 0.0355 | 0.205 | 1.449771647 |
| Nicotinate and Nicotinamide Metabolism | 0.043 | 0.219 | 1.366531544 |
| Aspartate Metabolism | 0.043 | 0.219 | 1.366531544 |
| De Novo Triacylglycerol Biosynthesis | 0.0447 | 0.219 | 1.349692477 |
| Carnitine Synthesis | 0.0503 | 0.224 | 1.298432015 |
| Transfer of Acetyl Groups into Mitochondria | 0.0503 | 0.224 | 1.298432015 |
| Glycolysis | 0.0563 | 0.24 | 1.249491605 |
| Pyrimidine Metabolism | 0.0649 | 0.255 | 1.187755303 |
| Glycerol Phosphate Shuttle | 0.0651 | 0.255 | 1.186419011 |
| Propanoate Metabolism | 0.0756 | 0.267 | 1.121478204 |
| Phosphatidylethanolamine Biosynthesis | 0.0762 | 0.267 | 1.118045029 |
| Cysteine Metabolism | 0.0763 | 0.267 | 1.117475462 |
| Ketone Body Metabolism | 0.0879 | 0.297 | 1.056011125 |
| Pyruvate Metabolism | 0.105 | 0.339 | 0.978810701 |
| Inositol Metabolism | 0.107 | 0.339 | 0.970616222 |
| Starch and Sucrose Metabolism | 0.116 | 0.354 | 0.935542011 |
| Methylhistidine Metabolism | 0.147 | 0.437 | 0.832682665 |
| Butyrate Metabolism | 0.167 | 0.466 | 0.777283529 |
| Galactose Metabolism | 0.18 | 0.466 | 0.744727495 |
| Nucleotide Sugars Metabolism | 0.181 | 0.466 | 0.742321425 |
| Glutathione Metabolism | 0.181 | 0.466 | 0.742321425 |
| Threonine and 2-Oxobutanoate Degradation | 0.181 | 0.466 | 0.742321425 |
| Tryptophan Metabolism | 0.192 | 0.483 | 0.716698771 |
| Methionine Metabolism | 0.221 | 0.529 | 0.655607726 |
| Histidine Metabolism | 0.221 | 0.529 | 0.655607726 |
| Inositol Phosphate Metabolism | 0.239 | 0.558 | 0.621602099 |

| | | | |
|---|---|---|---|
| Glycerolipid Metabolism | 0.254 | 0.578 | 0.595166283 |
| Phytanic Acid Peroxisomal Oxidation | 0.269 | 0.585 | 0.57024772 |
| Plasmalogen Synthesis | 0.269 | 0.585 | 0.57024772 |
| Phenylalanine and Tyrosine Metabolism | 0.283 | 0.59 | 0.548213564 |
| Selenoamino Acid Metabolism | 0.283 | 0.59 | 0.548213564 |
| Phenylacetate Metabolism | 0.301 | 0.59 | 0.521433504 |
| Homocysteine Degradation | 0.301 | 0.59 | 0.521433504 |
| Phospholipid Biosynthesis | 0.313 | 0.59 | 0.504455662 |
| Pentose Phosphate Pathway | 0.313 | 0.59 | 0.504455662 |
| Folate Metabolism | 0.313 | 0.59 | 0.504455662 |
| Lysine Degradation | 0.328 | 0.597 | 0.484126156 |
| Pyruvaldehyde Degradation | 0.329 | 0.597 | 0.482804102 |
| Fructose and Mannose Degradation | 0.342 | 0.61 | 0.465973894 |
| Phosphatidylcholine Biosynthesis | 0.429 | 0.75 | 0.367542708 |
| Porphyrin Metabolism | 0.468 | 0.805 | 0.329754147 |
| Phosphatidylinositol Phosphate Metabolism | 0.494 | 0.834 | 0.306273051 |
| Tyrosine Metabolism | 0.522 | 0.856 | 0.282329497 |
| Lactose Synthesis | 0.533 | 0.856 | 0.273272791 |
| Ethanol Degradation | 0.533 | 0.856 | 0.273272791 |
| Pantothenate and CoA Biosynthesis | 0.569 | 0.886 | 0.244887734 |
| Betaine Metabolism | 0.569 | 0.886 | 0.244887734 |
| Caffeine Metabolism | 0.603 | 0.919 | 0.219682688 |
| Androstenedione Metabolism | 0.619 | 0.919 | 0.208309351 |
| Estrone Metabolism | 0.619 | 0.919 | 0.208309351 |
| Oxidation of Branched Chain Fatty Acids | 0.649 | 0.933 | 0.187755303 |
| Mitochondrial Beta-Oxidation of Short Chain Saturated Fatty Acids | 0.663 | 0.933 | 0.178486472 |
| Mitochondrial Beta-Oxidation of Medium Chain Saturated Fatty Ac | 0.663 | 0.933 | 0.178486472 |
| Pterine Biosynthesis | 0.676 | 0.933 | 0.170053304 |
| Mitochondrial Beta-Oxidation of Long Chain Saturated Fatty Acids | 0.676 | 0.933 | 0.170053304 |
| Bile Acid Biosynthesis | 0.735 | 0.988 | 0.133712661 |
| Androgen and Estrogen Metabolism | 0.736 | 0.988 | 0.133122186 |
| Fatty Acid Elongation In Mitochondria | 0.757 | 0.989 | 0.12090412 |
| Retinol Metabolism | 0.757 | 0.989 | 0.12090412 |
| Sphingolipid Metabolism | 0.802 | 1 | 0.095825632 |
| Fatty acid Metabolism | 0.825 | 1 | 0.083546051 |
| Steroidogenesis | 0.825 | 1 | 0.083546051 |
| Steroid Biosynthesis | 0.858 | 1 | 0.066512712 |
| Arachidonic Acid Metabolism | 0.936 | 1 | 0.028724151 |

| Produced by L. crispatus B only | | | |
|---|---|---|---|
| **Metabolic pathways** | **Raw p** | **FDR** | **minus log 10 p** |
| Glycine and Serine Metabolism | 0.000603 | 0.0591 | 3.219682688 |
| Betaine Metabolism | 0.0168 | 0.823 | 1.774690718 |
| Spermidine and Spermine Biosynthesis | 0.0821 | 1 | 1.085656843 |
| Catecholamine Biosynthesis | 0.0986 | 1 | 1.006123085 |
| Glutathione Metabolism | 0.0986 | 1 | 1.006123085 |
| Methionine Metabolism | 0.0996 | 1 | 1.001740662 |
| Inositol Phosphate Metabolism | 0.134 | 1 | 0.872895202 |
| Urea Cycle | 0.172 | 1 | 0.764471553 |
| Inositol Metabolism | 0.192 | 1 | 0.716698771 |
| Homocysteine Degradation | 0.219 | 1 | 0.659555885 |
| Nicotinate and Nicotinamide Metabolism | 0.242 | 1 | 0.616184634 |
| D-Arginine and D-Ornithine Metabolism | 0.261 | 1 | 0.583359493 |
| Galactose Metabolism | 0.273 | 1 | 0.563837353 |
| Taurine and Hypotaurine Metabolism | 0.281 | 1 | 0.55129368 |
| Phosphatidylethanolamine Biosynthesis | 0.281 | 1 | 0.55129368 |
| Glucose-Alanine Cycle | 0.3 | 1 | 0.522878745 |
| Phosphatidylcholine Biosynthesis | 0.32 | 1 | 0.494850022 |
| Alanine Metabolism | 0.374 | 1 | 0.427128398 |
| Phosphatidylinositol Phosphate Metabolisn | 0.374 | 1 | 0.427128398 |
| Warburg Effect | 0.462 | 1 | 0.335358024 |
| Glycolysis | 0.47 | 1 | 0.327902142 |
| Valine, Leucine and Isoleucine Degradation | 0.48 | 1 | 0.318758763 |
| Tryptophan Metabolism | 0.48 | 1 | 0.318758763 |
| Glycerolipid Metabolism | 0.499 | 1 | 0.301899454 |
| Phenylalanine and Tyrosine Metabolism | 0.526 | 1 | 0.279014256 |
| Selenoamino Acid Metabolism | 0.526 | 1 | 0.279014256 |
| Bile Acid Biosynthesis | 0.533 | 1 | 0.273272791 |
| Phospholipid Biosynthesis | 0.552 | 1 | 0.258060922 |
| Lysine Degradation | 0.565 | 1 | 0.247951552 |
| Tyrosine Metabolism | 0.575 | 1 | 0.240332155 |
| Starch and Sucrose Metabolism | 0.577 | 1 | 0.238824187 |
| Fructose and Mannose Degradation | 0.577 | 1 | 0.238824187 |
| Citric Acid Cycle | 0.589 | 1 | 0.229884705 |
| Amino Sugar Metabolism | 0.6 | 1 | 0.22184875 |
| Gluconeogenesis | 0.6 | 1 | 0.22184875 |
| Beta-Alanine Metabolism | 0.611 | 1 | 0.21395879 |
| Sphingolipid Metabolism | 0.672 | 1 | 0.172630727 |
| Glutamate Metabolism | 0.739 | 1 | 0.131355562 |
| Arginine and Proline Metabolism | 0.767 | 1 | 0.115204636 |
| Pyrimidine Metabolism | 0.799 | 1 | 0.097453221 |

| Produced by L. crispatus S only | | | |
|---|---|---|---|
| **Metabolic pathways** | **Raw p** | **FDR** | **minus log 10 p** |
| Purine metabolism | 0.000252 | 0.0211 | 3.598599459 |
| Phenylalanine metabolism | 0.00383 | 0.159 | 2.416801226 |
| Tyrosine metabolism | 0.00688 | 0.159 | 2.162411562 |
| Arginine biosynthesis | 0.00757 | 0.159 | 2.12090412 |
| Histidine metabolism | 0.00987 | 0.166 | 2.005682847 |
| Citrate cycle (TCA cycle) | 0.0153 | 0.214 | 1.815308569 |
| Alanine, aspartate and glutamate metabolism | 0.0291 | 0.349 | 1.536107011 |
| Phenylalanine, tyrosine and tryptophan biosynthesis | 0.0385 | 0.405 | 1.41453927 |
| D-Glutamine and D-glutamate metabolism | 0.0573 | 0.492 | 1.241845378 |
| Tryptophan metabolism | 0.0586 | 0.492 | 1.232102384 |
| Ubiquinone and other terpenoid-quinone biosynthes | 0.0847 | 0.647 | 1.07211659 |
| Butanoate metabolism | 0.137 | 0.962 | 0.863279433 |
| beta-Alanine metabolism | 0.187 | 1 | 0.728158393 |
| Pyruvate metabolism | 0.195 | 1 | 0.709965389 |
| Pyrimidine metabolism | 0.321 | 1 | 0.493494968 |
| Aminoacyl-tRNA biosynthesis | 0.38 | 1 | 0.420216403 |
